# Genomic Architecture, Differentiation, and Adaptation in Three Large Falcons

**DOI:** 10.64898/2026.05.21.726861

**Authors:** Justin J. S. Wilcox, Barbara Arca-Ruibal, Stéphane Boissinot, Youssef Idagdour

## Abstract

Recent chromosomal rearrangements and divergence in large falcon species make them excellent foci for studies on evolution and genomic architecture. Here, we use high-coverage (44-74X) whole genome resequencing with 10X Genomics Linked-Reads to assess patterns of genomic divergence in peregrine, saker, and gyrfalcons and we link these to chromosomal type and chromosomal rearrangements. We first use admixture analysis and cross-coalescent MSMC2 to demonstrate distinct species boundaries between the large falcons and retrace their demography. We assessed genomic landscapes in terms of recombination rate, nucleotide diversity (π), Tajima’s D, autozygosity and F_st_ between saker and gyrfalcons: π had higher values on smaller chromosomes and F_st_ had higher values on larger chromosomes. Recombination rate concealed other chromosome type effects on π and Tajima’s D but largely explained variation in F_st_. We find 39 selective sweeps—some shared—across the falcons. However, only five candidate genes—mostly housekeeping genes—were implicated as targets of balancing selection across all falcons, with 4 of these shared between *Hierofalco* and three shared across all the falcons. Occurrence of selective sweeps and balancing selection were not enriched by chromosome type or in the context of chromosome fusions. Overall, our findings provide insights into divergence and adaptation in large falcons, and demonstrate an association of genomic architecture and chromosomal fusions with all population genomic indicators and metrics of differentiation between species.

**Significance Statement:** Falcons are culturally and commercially important birds that have undergone recent chromosomal rearrangement, providing a natural system for studies on chromosomal heterogeneities and evolution. By analyzing genomic variation across three large falcon species, we show that chromosome type and chromosomal fusions structure patterns of recombination, diversity, and divergence. Our findings highlight the importance of underlying genomic architecture to common forms of evolutionary inference and call attention to the role of chromosomal fusions in shaping falcon evolution.

## Introduction

Speciation is a continuum of reproductive isolation often conceptualized as resulting from divergent selective forces (Stankowski & Ravinet, 2021). Under this classical paradigm, mutations randomly arise within a genome and are subjected to varying degrees of drift and selection depending on the size of, phenotypic impacts on, and selective landscape experienced by the population(s) in which they occur. Following sufficient genomic divergence, selective pressures against interbreeding can arise. Though research efforts have historically focused on the extrinsic roles of the geographic landscape and the ecological niche on divergence, adaptation and speciation, genomes are themselves highly heterogenous, and heterogeneities in the genomic landscape have the capacity to influence all aspects of this evolutionary process (Dauphin et al., 2023; Storfer et al., 2018). Mutations do not occur or fix equally across the genome (Francioli et al., 2015; Mugal et al., 2015; Nesta et al., 2021), and differences in recombination rates effectively alter the effective population size of each genomic region (Charlesworth, 2009). Regions with higher mutation rate should have more genetic variability. Regions with higher recombination rate should have reduced background selection resulting in more variation, higher effective population sizes, slower rates of differentiation, and faster fixation of beneficial variation due to reduced Hills-Robertson Interference. As such, the impacts of drift, selection, and introgression are likely to differ across a genome. Though well established in theory and comparative genomics (Axelsson et al., 2005; Ellegren, 2013; Weir et al., 2005), the practical importance of these within-genome effects on evolution are of emerging interest (Cicconardi et al., 2021; Manthey et al., 2021; Mezzasalma et al., 2024; Nasvall et al., 2023; Palahi et al., 2023; Schield et al., 2020; Yoshida et al., 2023) but largely undescribed at present. Recent advances in sequencing technology now allow for the contiguous high-coverage genomes necessary to empirically explore these processes. These processes are likely to play a pronounced role in avian evolution, particularly in falcons.

Birds possess atypically heterogenous genomes—with extreme variation in chromosome size, recombination rate, GC content, CpG islands density, and gene density (Ellegren, 2013)—and are among the most species rich groups of amniotes (Callaghan et al., 2021). These heterogeneities are intricately linked to a conserved avian genomic organization consisting of macrochromosomes (40-200+ MB), intermediate chromosomes (20-40 MB), and microchromosomes (1-20 MB). Though most of the DNA in a typical avian genome is carried on the macrochromosomes, most of the genes are located on microchromosomes, which also display higher mutation rates, recombination rate, GC content, and counts of CpG sites and islands (Ellegren, 2010; Wilcox et al., 2022). While this organizational structure is believed to be plesiomorphic to all amniotes it has generally been lost or reduced in most constituent clades other than birds (Waters et al., 2021). This parallels an unusually conserved organization of avian karyotypes, such that the majority of birds possess 2N∼82 (Ellegren, 2010; O’Connor, Romanov, et al., 2018). While the evolutionary significance of this karyotype and of the broader genomic organizational structure of birds are unknown, their conserved nature would suggest that they are important to the long-term survival of avian lineages. This organizational structure has also been shown to drive differential population genomic signatures across avian genomes with larger chromosomes appearing to accelerate lineage sorting during speciation (Manthey et al., 2021; Manthey et al., 2016). It is unclear, however, if this is strictly the result of differences in recombination rate or other aspects of macrochromosomes due to the many differences between avian chromosome types. The many unique traits of microchromosomes—particularly high gene content, hypermutability, and enrichment for epigenetically active sites—may likewise play other important roles in neutral and adaptive evolution of birds, though this remains generally unassessed. The many peculiar features of falcon genomes make them excellent foci for studies on avian evolution (Wilcox et al., 2022).

Falcons (Genus: *Falco*) contain the fastest and most broadly distributed species of bird, the peregrine falcon, and have undergone several radiations in the last ∼7.5 million years (Fuchs et al., 2015; Wilcox et al., 2019). The diversity and ecological peculiarities of falcons have arisen in the context of an unusual genomic architecture, as falcons have undergone a series of chromosomal rearrangements—consisting primarily of fusions of microchromosomes to macrochromosomes—to arrive at a karyotype of 2N of 50-52 (Joseph, 2017; O’Connor, Farre, et al., 2018). These chromosomal fusions provide a natural experiment to disentangle the otherwise highly correlated traits of avian genomic organization (Joseph et al., 2018; O’Connor, Farre, et al., 2018; Wilcox et al., 2022), and have thrown falcon genomes out of AT-GC equilibrium, driven a major loss of GC content and CpG sites, and affected almost all other aspects of genomic architecture from NUMT insertion rates to transposable element (TE) clearance (Wilcox et al., 2022). The population genomic consequences of these chromosomal fusions remain unexplored, however, as do their potential roles in driving the unique adaptive potential of falcons. These ecological and genomic peculiarities, nonetheless, pair with extensive prior sequencing of falcons (Cho et al., 2019; Johnson et al., 2023; Joseph et al., 2018; O’Connor, Farre, et al., 2018; Wilcox et al., 2022; Zhan et al., 2013; Zuccolo et al., 2023) to make falcons prime targets for studies on avian evolution in the context of genomic architecture.

Falcons can be broken into 10 major clades consisting of 39 species in total (Fuchs et al., 2015; Wilcox et al., 2019). With 10 species, large falcons are among the most species rich of these clades and diverged from other falcons in the early Pleistocene ∼2.5 million years ago. This clade contains the subgenus *Hierofalco*, which consists of four species that have diverged in the last 300-400 KYA and occur across Eurasia and Africa, and the peregrine-species complex, which as the single most broadly-distributed species of bird contains 19 (currently-recognized) subspecies across all major land masses with the notable exception of Antarctica (Wilcox et al., 2019). *Hierofalco* and peregrine falcons are sister clades that diverged from one another within the last half-million to one-million years (Fuchs et al., 2015; Wilcox et al., 2022). These clades generally correspond to different ecological strategies and phenotypes: peregrine falcons occupy highly-diverse habitats (Johnson et al., 2023; Wink, 2018) but specialize on avian prey (Cresswell, 1996; Dekker, 1988; Jenkins, 1995, 2000)—with the unique and notable capacity to exceed 300 Km/hr in a stoop or dive (Ponitz et al., 2014)—whereas *Hierofalco* spp. tend to be specialized in their habitats but generalists on mammalian, avian, and reptilian prey (Chavko et al., 2014; Gombobaatar et al., 2001; Poole & Boag, 1988; Poole & Bromley, 1988; White & Weeden, 1966). Due to this habitat specialization, *Hierofalco* spp. tend to be allopatric or parapatric to one another and natural hybridization is likewise not known to be common (Wink et al., 2004). The arctic-adapted gyrfalcon and arid-adapted saker falcon are the most recently diverged species of *Hierofalco* with a divergence time within the last 20-300 thousand years (Al-Ajli et al., 2023; Hu et al., 2022; Pan et al., 2017; Wilcox et al., 2022). Hybridization between the species is considerably rare but possible (Moseikin & Ellis, 2004), though strong empirical support for this is lacking. Ultimately, peregrine falcons, gyrfalcons, and saker falcons have all recently evolved and adapted under differential selective pressures. These differences and the lack of post-zygotic isolation between these species make their divergence times, processes of speciation, and demographic histories of particular interest (Wilcox et al., 2019; Zinevich et al., 2023).

Here we use 10X Genomics Chromium linked-read sequencing (Marks et al., 2019) to characterize the population genomics of peregrine falcons (N=11), gyrfalcons (N=14), and saker falcons (N=10) in the context of genomic architecture and chromosomal fusions. We confirm that the falcons represent three distinct species, and use within species coalescents to show distinct demographic trajectories for each falcon. Using cross-species coalescents, we find that peregrine falcons were reproductively isolated for hundreds of thousands of years, but have had more recent gene flow with both species of *Hierofalco*. We also find that saker and gyrfalcons had reducing geneflow over approximately 400,000 years, with apparent reproductive isolation occurring only approximately 12,000 years ago. Prevailing theory has historically explained genomic heterogeneities primarily in the context of recombination and would predict lower π, Tajima’s D, and Fst on larger chromosomes due to lower recombination and increased background selection. However, the many related heterogeneities related to chromosome type (Ellegren, 2013) suggest that chromosome-size specific patterns of evolution should often occur independently of recombination rate. We confirm that fusions of smaller chromosomes onto larger chromosomes lead to reductions in recombination rate, nucleotide diversity (π), Tajima’s D and lineage sorting between species. We also find that recombination rate, π, decrease with chromosome size and that F_st_ between species increases with chromosome size. However, differences in recombination rate conceal broader patterns of π and Tajima’s D that are associated with chromosome type. While regional differences in F_st_ between peregrine falcons and *Hierofalco* appear to be driven by recombination, differences in recombination rate conceal chromosome-type-specific differences in F_st_ between gyr and saker falcons as well. We find evidence for limited selective sweeps and balancing selection on the falcon genomes, but find no association between selective sweeps and chromosome type or chromosomal fusions.

## Results

### Resequencing and Alignment

Illumina Novaseq sequencing was performed on high-molecular-weight DNA extracted from 35 blood samples collected from 14 gyrfalcons, 11 peregrine falcons, and 10 saker falcons and prepared using 10X Chromium linked-reads library preparation. Demultiplexing produced 19,385,467,990 barcoded paired-end reads with an average of 553,870,514 per sample. Reads were aligned to a scaffold-scale lanner falcon reference genome, GCA_024433245.1 (Wilcox et al., 2022), using 10X Genomics Longranger software, which takes linked-read information into account. Species-specific effects in sequencing, alignment, and phasing efficacy were observed (Supplementary Figure 1-5). Mean depth averaged 58X (range 44X-74X), with an average coverage of 99.9%, >99% phasing success for all samples, and a minimum phaseblock N50 of 1,466,213 with many samples phasing far better (mean= 3522893; maximum= 6,458,353).

### Summary Statistics

Allele frequency spectra were produced for each falcon species using single-nucleotide variants (SNVs), consisting of all single-base substitutions regardless of frequency (Fig. 1A). Distributions differed significantly between all species (Kruskall-Wallace Test: Kruskal–Wallis χ² = 124,836; DF=2; p<10^-6^; ε²=0.012111; R^2^=0.01232. Pairwise Mann-Whitney-U tests: p<10^-6^ for all. Pairwise Kolmogorv-Smirnov Tests: Gyr-Saker D=0.23178; Gyr-Peregrine D=0.23178; Saker-Peregrine D=0.33444; p<10^-6^ for all). However, all curves had similar shapes with rare variants being most common but a slight excess of more common variants, suggesting that the lanner falcon outgroup is of insufficient distance to polarize all variant calls. Gyrfalcons also had a lower level of singleton variants relative to the other falcon species. Genome-wide patterns of Tajima’s D were slightly negative for all falcons (Fig. 1B; Supplementary Figure 6), but differed in extent between all species (Kruskal-Wallace: χ^2^= 4,005.6; DF=2; p<10^-6^; ε²= 0.12416; R^2^=0.1299. Pairwise Mann-Whitney-U tests: p<10^-6^ for all). Recombination rate (Fig. 1 C), r, demonstrated significantly different distributions between all species (Kruskal-Wallace: χ^2^=7,993.3; DF=2; p<10^-6^; ε²= 0.24947; R^2^=0.1911. Pairwise Mann-Whitney-U tests: p<10^-6^ for all), but several distinct regions with elevated diversity and depressed recombination rate occurred in all species (Supplementary Figure 7). Among 100KB windows, recombination rate showed a skewed distribution in all species with the highest mean recombination rate in peregrine falcon and lowest mean recombination rate within the gyrfalcon. Genomic windows of nucleotide diversity (π) were similar across species (Fig. 1D), and resembled patterns seen in recombination rate, with several shared regions of elevated diversity (Supplementary Figure 8). Distributions were still significantly different between species (Kruskal-Wallace: χ^2^= 9,978.4; DF=2; p<10^-6^; ε²= 0.30956; R^2^=0.2186. Pairwise Mann-Whitney-U tests: p<10^-6^ for all) with the highest diversity in the peregrine falcon and the lowest diversity in gyrfalcon.

**Figure 1:**
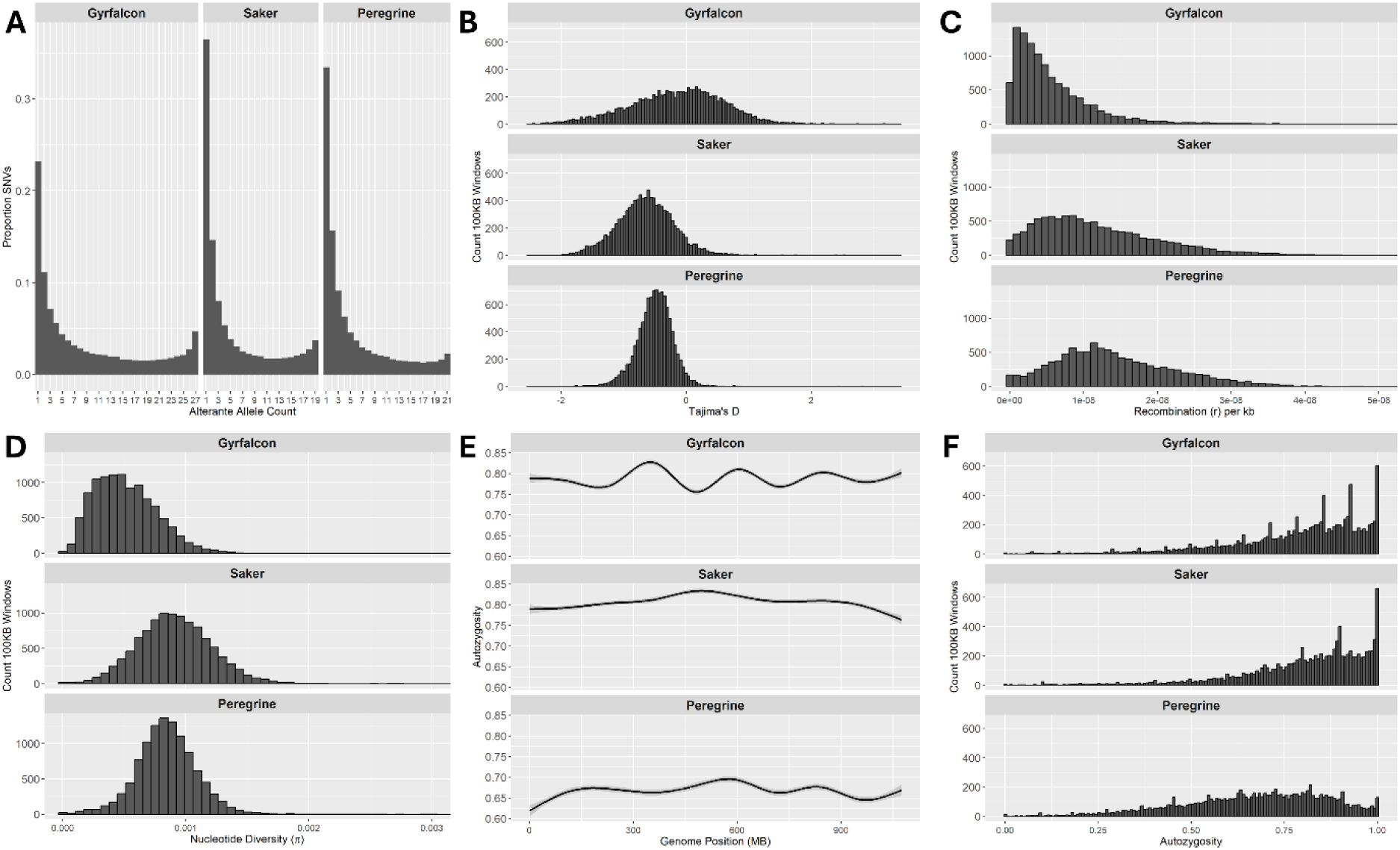
Summary statistics on within species population genomics for each falcon species. A) Allele-frequency spectra; B) Distribution of Tajima’s D across 100KB windows; C) Distribution of recombination rate per base (r) across 100KB windows; D) Distribution of π across 100KB windows; E) Autozygosity across each genome; F) Distribution of autozygosity across 100KB windows.

Distributions of autozygosity differed between all species (Fig. 1E & 1F), albeit with low effect size (Kruskal-Wallace: χ^2^=9,418.3; DF=2; p<10^-6^; ε²= 0.02504; R^2^=0.025. Pairwise Mann-Whitney-U tests: p<10^-6^ for all). Shared autozygosity was very rare within species but, autozygosity was common among individuals. The overwhelming majority (Gyrfalcon: 99.5%; Saker: 98.8; Peregrine: 98.2%) of 100kb windows were autozygous in at least one individual of each species. However, shared runs of autozygosity of 100kb (F_roh(100kb)_) were proportionally low and accounted for 4.12% of the gyrfalcon genome, 4.48% of the saker falcon genome, and 0.95% of the peregrine falcon genome. The longest shared run of autozygosity in the gyrfalcon was between 300-400KB. The longest shared runs of autozygosity in the saker and peregrine falcons were 200-300KB. No runs of autozygosity were shared between all species or within the subgenus *Hierofalco*.

### Admixture and Demography

We examined broad demographic patterns of differentiation between species. Unique and shared SNVs and indels were calculated across autosomal scaffolds of each species with CpG sites and non-CpG sites separated (Fig. 2A). Admixture analysis of 100,000 randomly sampled SNVs showed no admixture between species (Fig. 2B).

**Figure 2:**
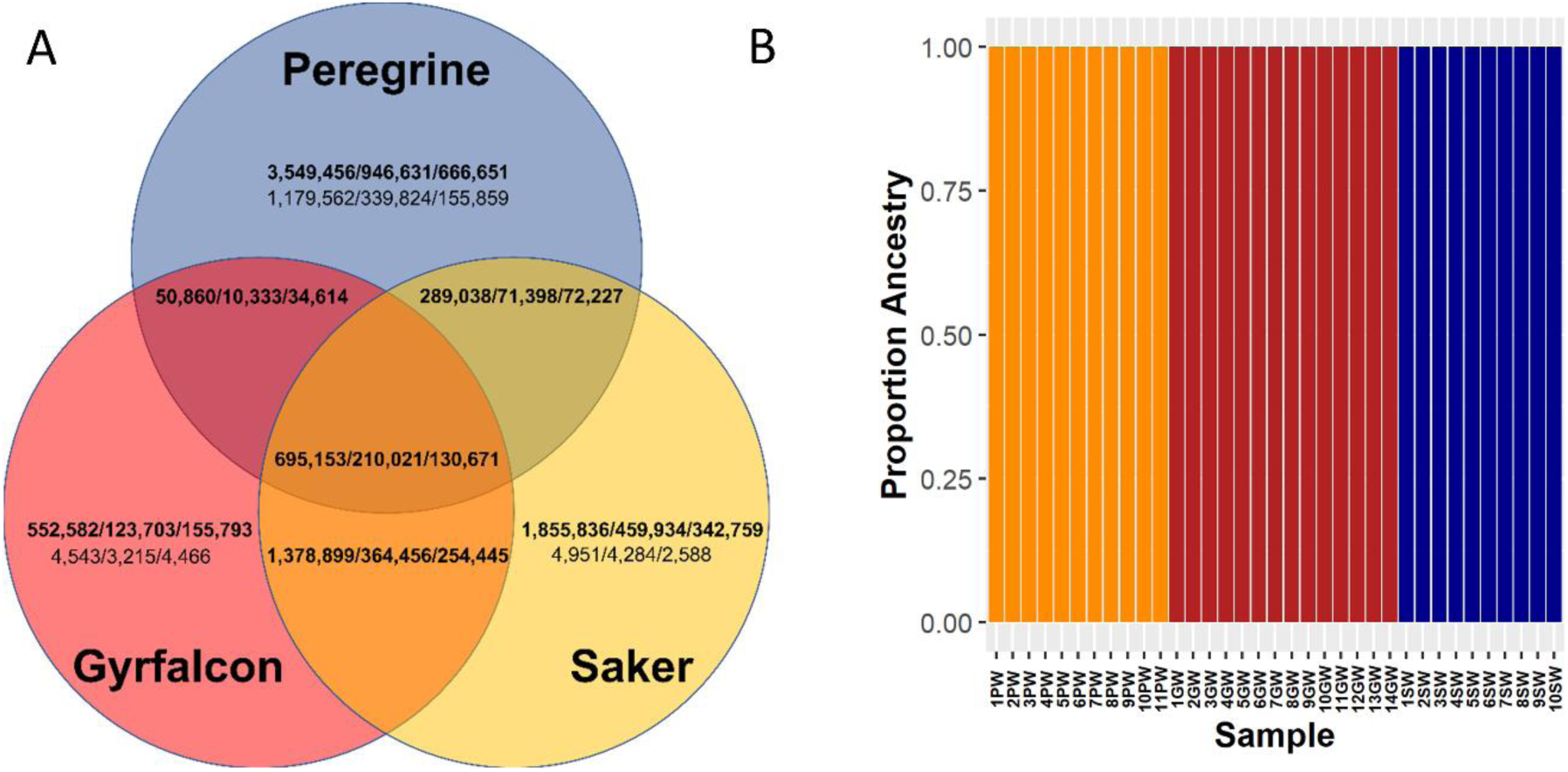
Shared and unique diversity with target falcons. A) Venn diagram of small variants by type and context (SNVs-Outside-CpG-Context/SNVs-Within-CpG-Context/Indels) shared within and between falcon species. Fixed differences between groups are shown in non-bold text. B) Admixture plot based on 100,000 subsampled SNVs from each falcon species.

Variants types differed in their levels of differentiation within and between species. SNVs at non-CpG sites were less likely to be shared between species (Fisher’s Exact Test Peregrine: Odds Ratio=0.9650967 ; p<10^-6^; Fisher’s Exact Test Gyrfalcon: Odds Ratio=0.8277268; p<10^-6^; Fisher’s Exact Test Saker: Odds Ratio=0.9127781; p<10^-6^), and more likely to be polymorphic within species than SNVs at CpG sites in all falcon species (Fisher’s Exact Test Peregrine: Odds Ratio=1.080222; p<10^-6^; Fisher’s Exact Test Gyrfalcon: Odds Ratio=3.161316; p<10^-6^; Fisher’s Exact Test Saker: Odds Ratio=3.491434; p<10^-6^)—that is, SNV variation at CpG sites was more likely to be private to and fixed within falcon species. SNV variation showed a less decisive pattern relative to indels. When CpG sites are excluded, SNVs were more likely to be shared in gyrfalcons (Fisher’s Exact Test: Odds Ratio=1.456265; p<10^-6^), but less likely to be shared in peregrine (Fisher’s Exact Test: Odds Ratio= 0.7579604; p<10^-6^) and saker falcons (Fisher’s Exact Test: Odds Ratio=0.9589432; p<10^-6^), relative to indels. However, SNVs that were unique to saker (Fisher’s Exact Test: Odds Ratio= 2.830377; p<10^-6^) and gyrfalcons (Fisher’s Exact Test: Odds Ratio= 3.486567; p<10^-6^) were more likely to be polymorphic than indels, and more likely to be fixed in peregrine falcons (Fisher’s Exact Test: Odds Ratio=0.7035167; p<10^-6^) relative to indels. In short, no clear patterns emerged as to whether indels or SNVs were better at distinguishing species, although single nucleotide variation at CpG sites was more uniquely species specific than variation at non-CpG sites.

We reconstructed past demography and patterns of gene flow within and between falcon species. MSMC2 was run on 100 sets of four different haploid subsamples from each species to estimate demographic histories and variation in convergence times across the individuals sampled (Fig. 3). Within (Fig. 3A) and across species (Fig. 3B) coalescents were calculated for each bootstrap. Outputs of within-species coalescents were scaled to population size with the published collared flycatcher mutation rate (Smeds et al., 2016) and scaled to time with current estimated generation times for each species (Wilcox et al., 2022). Cross-species coalescents were scaled as a proportion of inverse coalescent rate, λ, to get an estimate of proportion of gene flow. Changes in generation time over the course of each species demographic history likely prevent exact convergence, but the saker and peregrine falcon show similar demographic trajectories from about 600,000 to 1,000,000 or more years ago. In the case of saker falcons and gyrfalcons different demographic trajectories appear about 500-800 thousand-years-ago. This is accompanied by a steady reduction in gene flow between the species until approximately 12,000 years ago. Results do not indicate any meaningful gene flow between the species since this time. Peregrine falcons consistently coalesced approximately one-million years ago and showed clearly distinct patterns of demographic history beginning between 650,000 and 700,000 years ago. Cross-species coalescence indicates that peregrine falcons rapidly achieved reproductive isolation from the ancestors of extant *Hierofalco* approximately 700,000 to one-million years ago and maintained reproductive isolation until approximately 100,000 to 200,000 years ago. Given similar patterns of cross-coalescence between peregrines and both saker falcons and gyrfalcons this appears to have begun before these two species achieved reproductive isolation. Present proportions of cross-coalescent ancestry between peregrine falcons and *Hierofalco* appear to be highly variable between individuals sampled but are always at least moderate. Additionally estimates of between peregrines and saker falcons or their ancestors are generally higher than those between gyrfalcons and their ancestors.

**Figure 3:**
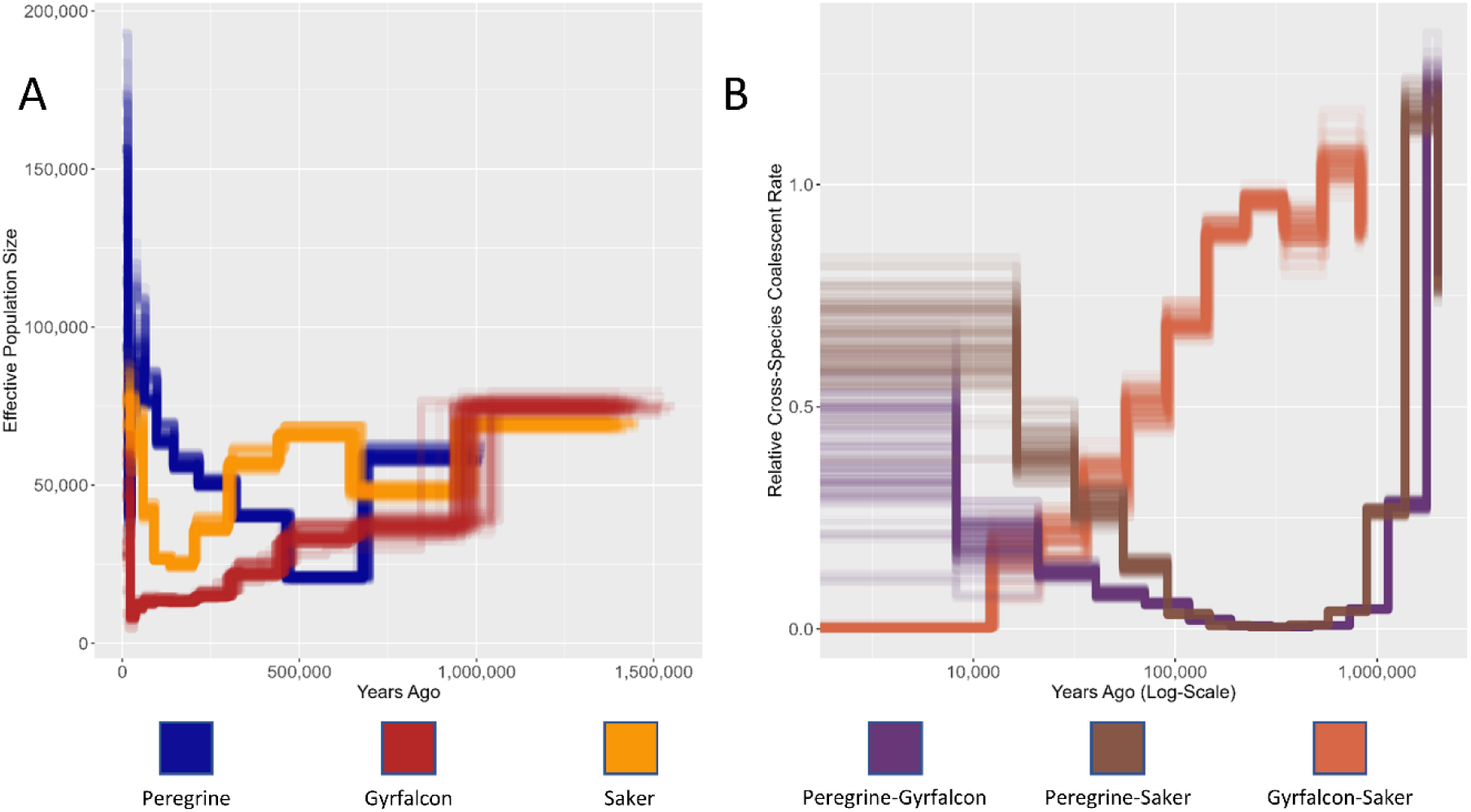
Within-species (A) and cross-species (B) MSMC2 for all falcon species. Each estimate is based on 100 subsampled bootstraps of 4 random falcon haplotypes for each species. Within-species and cross-species estimates are color-coded. Results are scaled to time and effective population size using reports of generation time for each falcon species and μ for the collared fly-catcher (Smeds et al., 2016). Cross-species coalescents are scaled to total coalescent within and between species (0-1 with 0 being reproductive isolation and 1 being panmixia). Time is estimated in cross-species coalescents using the harmonic mean of the generation time for each species compared and presented on a log_10_ scale.

Demographic trends since separation of peregrine falcons and the ancestors of *Hierofalco* show increasing peregrine falcon population sizes and decreasing *Hierofalco* population sizes. Saker falcon population sizes appear to stabilize and increase within the last 200,000 years. Gyrfalcon population sizes appear to begin increasing within the last few tens-of-thousands of years.

### Selection and Differentiation

We used allele frequency information and recombination rate to look for signatures of selection within falcon genomes. Scans for selective sweeps were performed with SweepFinder2. Scans revealed 19 genes that had undergone selective sweeps in the peregrine falcons, 12 in the gyrfalcons, and 11 in the saker falcon (Table 1). Some particular genes of interest stand out. The Tropomyosin-receptor kinase–fused gene (Protein TFG) appears to have undergone a selective sweep in all of the falcon species. This gene is involved in the function of the Golgi apparatus and is known to have impacts on metabolism, neural development, stress response, and immunity. Several other genes related to immunity also appear to be under directional selection in specific falcons, including Interleukin-8 (Gyrfalcons), CD166 antigen (Peregrine and saker), and Lysosomal-associated transmembrane protein 5 (Peregrine). All of these are involved in immune signaling and also link to effector functions in phagocytosis, cell adhesion, and turnover of immune cell surface receptors, respectively.

**Table 1:**
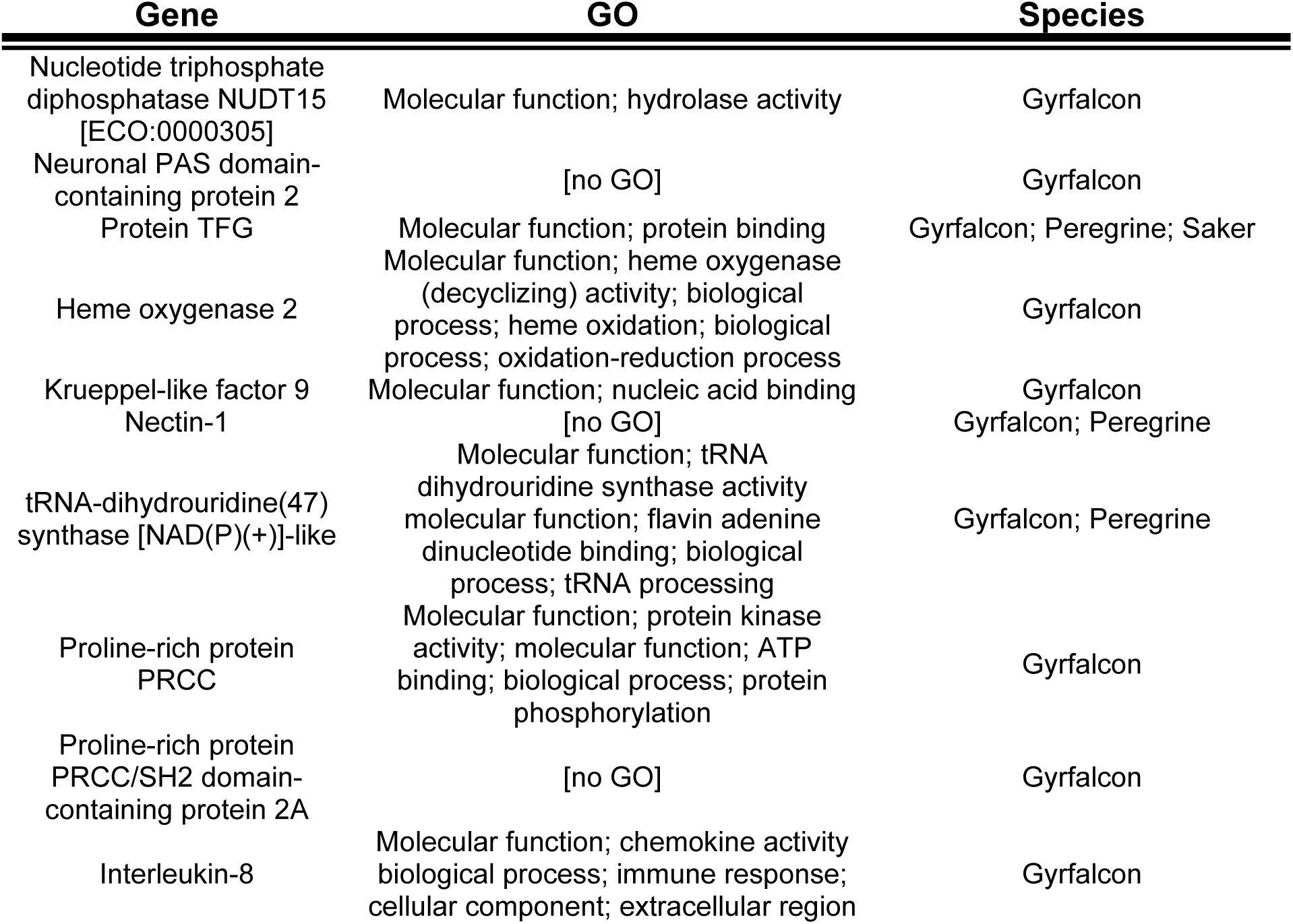
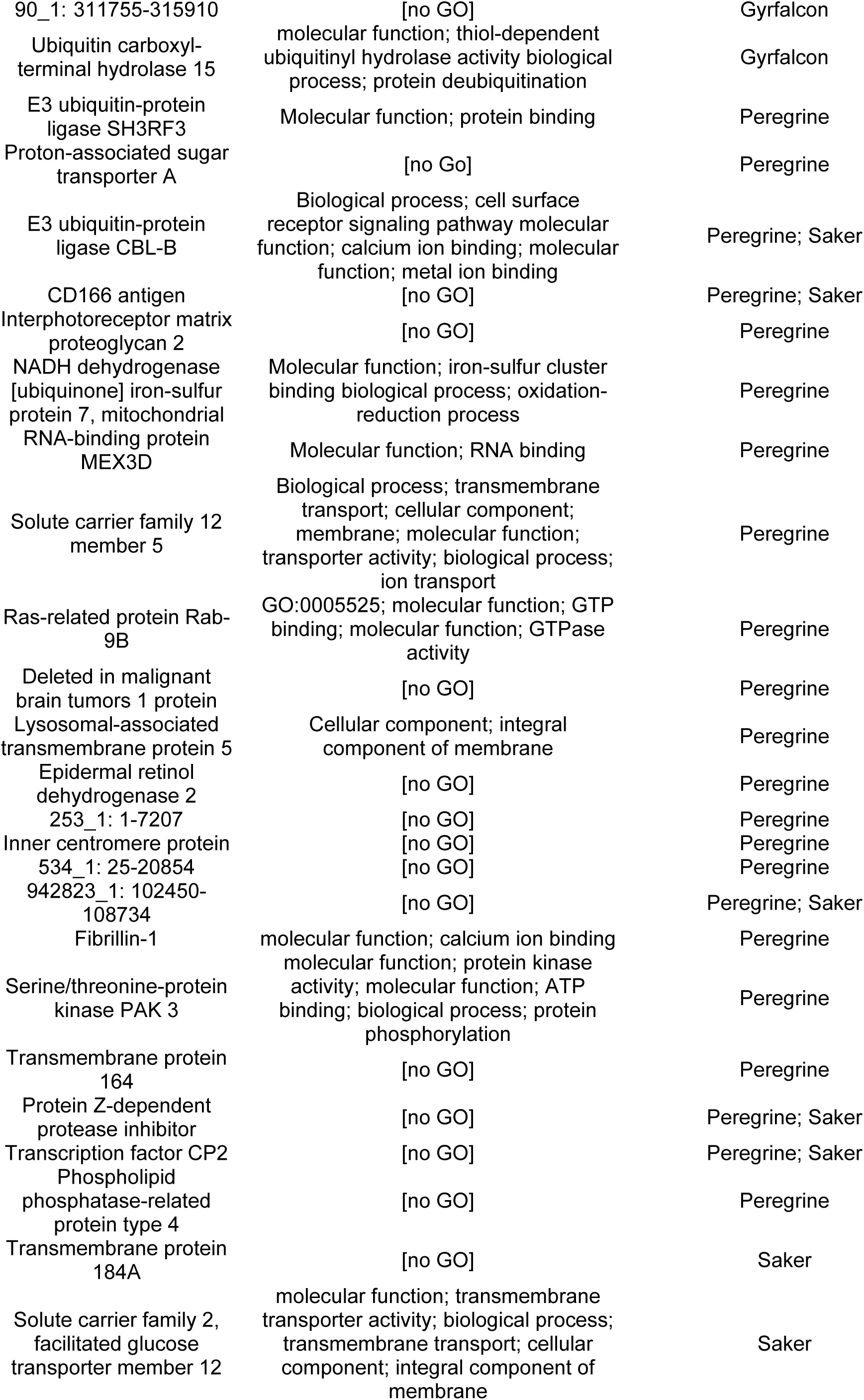
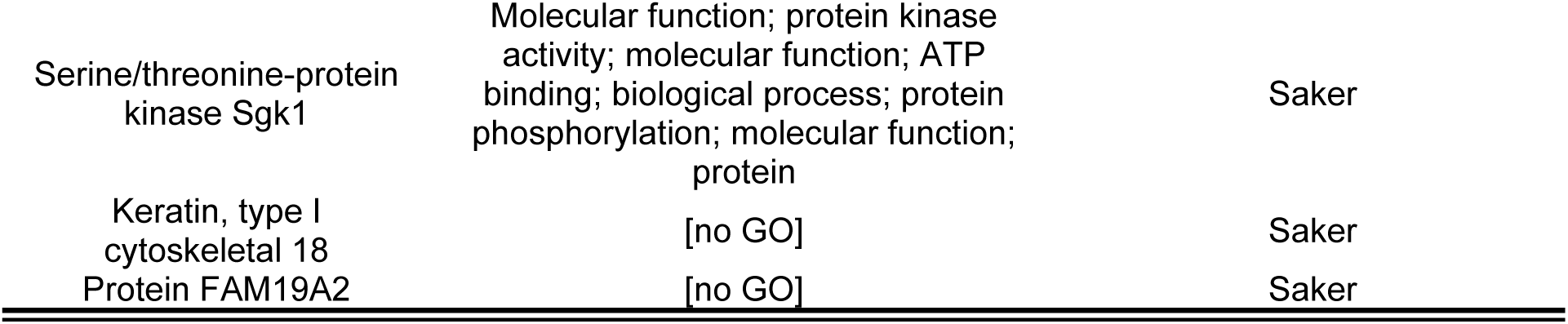
Genes Under Selective Sweeps in All Falcons. Unknown genes are listed by scaffold and coordinates.

Patterns of balancing selection are generally shared between falcons. Only five genes were found to be under balancing selection, four of which were shared between *Hierofalco* and three of which were shared between all three falcons. The one shared only between *Hierofalco* has no annotated functions but the rest have well known functions. One (E3 ubiquitin-protein ligase SMURF1), relates to ubiquitin signaling, particularly in immune function and cell differentiation. The next gene under balancing selection in all falcons was “Aspartate aminotransferase” which catalyzes the transformation of amino acids into other amino acids and is also used in aerobic respiration. Finally, the two remaining genes under balancing selection are for the same cytoplasmic actin in all three falcons, and a particular actin binding protein in only the gyrfalcon. Actin has a very wide range of functions including, cell development, mechanical performance, immune functions, and even gene expression.

### Influence of Genomic Architecture

We analyzed genome-wide heterogeneity across falcon species in the context of chromosomal configuration. Population genomic indicators were computed across aligned 100KB windows for all falcon species and assigned to a current chromosome type based on redundant alignment to chromosome-scale reference genomes for other falcons and ancestral chromosome type based on redundant alignments to the closest relatives of Falconidae (Wilcox et al., 2022), the songbirds, parrots, and seriemas—collectively the Australaves. Species-specific effects were regressed out and residuals averaged to assess overall associations of windows with chromosome type across species and additional post-hoc analyses were performed within each to assess potential species-specific deviations from averages. Results are reported for all current chromosome types—macrochromosomes (>40MB), intermediate chromosomes (20MB-40MB), and microchromosomes (<20MB)—when differences were seen between intermediate and macrochromosomes; when such differences are not observed results are reported for microchromosomes and large chromosomes (>20MB). Given the high degree of uncertainty in ancestral states, results are only reported for large chromosomes and microchromosomes.

We expected recombination rate to be higher on larger chromosomes and chromosomal fusions to reduce recombination rate. We find that distributions of recombination rates differed significantly (Kruskal-Wallace: χ^2^= 505.23; DF=2; p<10^-6^; ε²= 0.06118; R^2^=0.1367. Pairwise Mann-Whitney-U tests: p<10^-6^ for all) across all current chromosome types (Fig 4). Smaller chromosome types had generally higher recombination rates with microchromosomes tending toward higher recombination rates than intermediate chromosomes and macrochromosomes and intermediate chromosomes tending toward higher recombination rates than macrochromosomes. These patterns were replicated within each individual falcon species. When ancestral state is considered, distributions of recombination rate again differed significantly across all states (Kruskal-Wallace: χ^2^= 492.77; DF=2; p<10^-6^; ε²= 0.16700. Pairwise Mann-Whitney-U tests: p<10^-6^ for all; R^2^=0.3006) with regions conserved on microchromosomes tending toward higher recombination rates than those conserved on large chromosomes and windows formerly on microchromosomes and fused to macrochromosomes tending toward rates that were intermediate to both. These patterns were again replicated within each individual falcon species.

**Figure 4:**
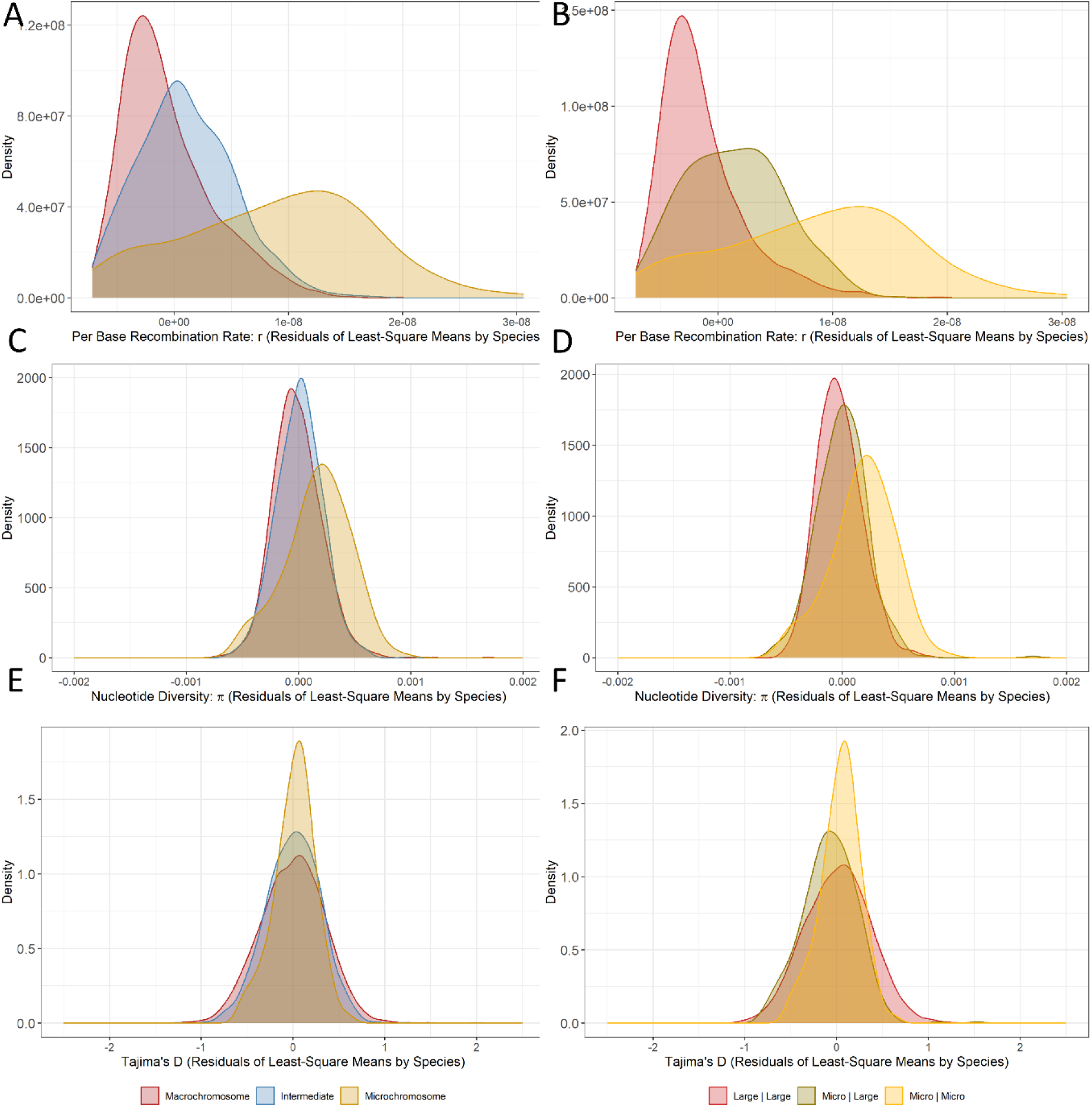
Recombination rate (r) and diversity (π) and Tajima’s D by current and ancestral chromosome. A) r by chromosome type; B) r by Ancestral | Current chromosome type. C) π by chromosome type; D) π by Ancestral | Current chromosome type. E) Tajima’s D by chromosome type; F) Tajima’s D by Ancestral | Current chromosome type.

We expected nucleotide diversity to be lower on larger chromosomes with a potential role for recombination rate in driving these trends. Distributions of π were weakly associated with chromosome type such that higher values π occurred on microchromosomes relative to other chromosome types (Fig. 4). Across all species (Kruskal-Wallace: χ^2^= 135.39; DF=2; p<10^-6^; ε²= 0.016217; R^2^=0.01317. Pairwise Mann-Whitney-U tests: p=0.00031 for macrochromosome-versus-intermediate; p<10^-6^ for others). Microchromosomes tended toward higher diversity than intermediate chromosomes and macrochromosomes, and intermediate chromosomes tended toward higher diversity than macrochromosomes. This pattern held for gyrfalcons, but intermediate chromosomes did not differ in diversity from macrochromosomes in saker and peregrine falcons. When we consider ancestral state (Kruskal-Wallace: χ^2^= 146.55; DF=2; p<10^-6^; ε²= 0.04922; R^2^=0.051. Pairwise Mann-Whitney-U tests: p<10^-6^ for Micro | Micro versus Micro | Large and Large | Large), we find that across all species windows conserved on microchromosomes tend toward higher π than those both conserved on and fused to large chromosomes from microchromosomes. These composite results hold within each individual species, and in the exclusive case of the gyrfalcon we also find that windows fused to large chromosomes from microchromosomes also tend toward higher π than regions conserved on large chromosomes (Pairwise Mann-Whitney-U tests: p=0.000006). Recombination rate was a strong predictor of π across falcon species (Linear Model: F=3,307; DF=8,226; p<10^-6^; R^2^=0.2867). Controlling for recombination rate by regressing out its effects led to a general reversal of sign (Supplementary Figure 9) such that larger chromosome types always tended toward higher π than expected based on their recombination rates (Kruskal-Wallace: χ^2^= 120.06; DF=2; p<10^-6^; ε²=0.014354; R^2^=0.01084. Pairwise Mann-Whitney U tests: p<10^-6^ for all). This trend held within each species although with the minor variation that microchromosomes and intermediate chromosomes did not differ in saker and peregrine falcons (Supplementary Figures 10-12). When ancestral state was considered (Supplementary Figure 13), π was again strongly correlated to recombination rate (Linear Model: F=1,656; DF=2,938; p<10^-6^; R^2^=0.3604). After controlling for recombination rate, π still weakly differed when ancestral state was considered (Kruskal-Wallace: χ^2^= 71.653; DF=2; p<10^-6^; ε²= 0.023715; R^2^= 0.02175. Pairwise Mann-Whitney U tests: p<10^-6^ for all except Micro | Micro versus Micro | Large), with microchromosomal fusions onto larger chromosomes tending toward lower values of π than expected based on recombination rate relative to regions conserved on large chromosomes. No differences were seen between regions conserved on microchromosomes and these fusions. However, within species analyses (Supplementary Figures 14-16) revealed these results to be entirely driven by patterns within saker and peregrine falcons (Pairwise Mann-Whitney U Tests: p<10^-6^ for fused regions versus regions conserved on large chromosomes in both species and fused regions versus those conserved on microchromosomes in peregrine falcons; p=0.00002 for fused regions versus regions conserved on microchromosomes in saker falcons) ; π did not significantly differ between fused and unfused regions within gyrfalcons when controlling for recombination rate.

We expected Tajima’s D to be lower on larger chromosomes as a result of reduced recombination. Distributions of Tajima’s D did not differ by current chromosome type either across all or within any falcon species (Fig. 5A & Fig. 5B). However, when ancestral state is considered (Kruskal-Wallace: χ^2^= 120.06; DF=2; p<10^-6^; ε²=0.012641; R^2^=0.01059. Pairwise Mann-Whitney U tests: p<10^-6^ for Micro | Large versus Micro | Micro and Large | Large), we find that windows that have fused to large chromosomes from microchromosomes have lower mean>Tajima’s D than those conserved on both large chromosomes and microchromosomes. However, these composite effects do not hold within individual species; fusions tended to have reduced Tajima’s D relative to only regions conserved on microchromosomes in gyrfalcons (Pairwise Mann-Whitney U tests: p<10^-6^) and relative to only regions conserved on large chromosomes (p=0.0000246) in saker and peregrine falcons (Pairwise Mann-Whitney U tests: p<10^-6^). Tajima’s D was correlated to recombination rate, but only weakly (Linear Model: F=548.6; DF=8,226; p<10^-6^; R^2^=0.06252). After regressing out the effects of recombination rate (Supplementary Figure 17), Tajima’s D differed significantly by current chromosome state with microchromosomes tending toward lower values of Tajima’s D than larger chromosomes (Pairwise Mann-Whitney U tests: p<10^-6^) across species. Within species (Supplementary Figures 18-20), gyrfalcons followed this general trend (Pairwise Mann-Whitney U tests: p<10^-6^). In saker falcons no significant difference was seen between microchromosomes and macrochromosomes but both microchromosomes (Pairwise Mann-Whitney U tests: p=0.000038) and intermediate (Pairwise Mann-Whitney U tests: p=0.00043) chromosomes tended toward lower values of Tajima’s D than macrochromosomes, and in peregrine falcons, smaller chromosome types always tended toward lower values of Tajima’s D than those that were larger (Pairwise Mann-Whitney U tests: p<10^-6^). When looking across (Supplementary Figure 21) and within each species (Supplementary Figures 22-24), regions of microchromosomes fused to larger chromosomes tended toward consistently lower values of Tajima’s D than those conserved on large chromosomes across and within all species (Pairwise Mann-Whitney U tests: p<10^-6^ for all species, saker falcon, and peregrine falcon; p=0.0000016 for gyrfalcons) after controlling for recombination.

**Figure 5:**
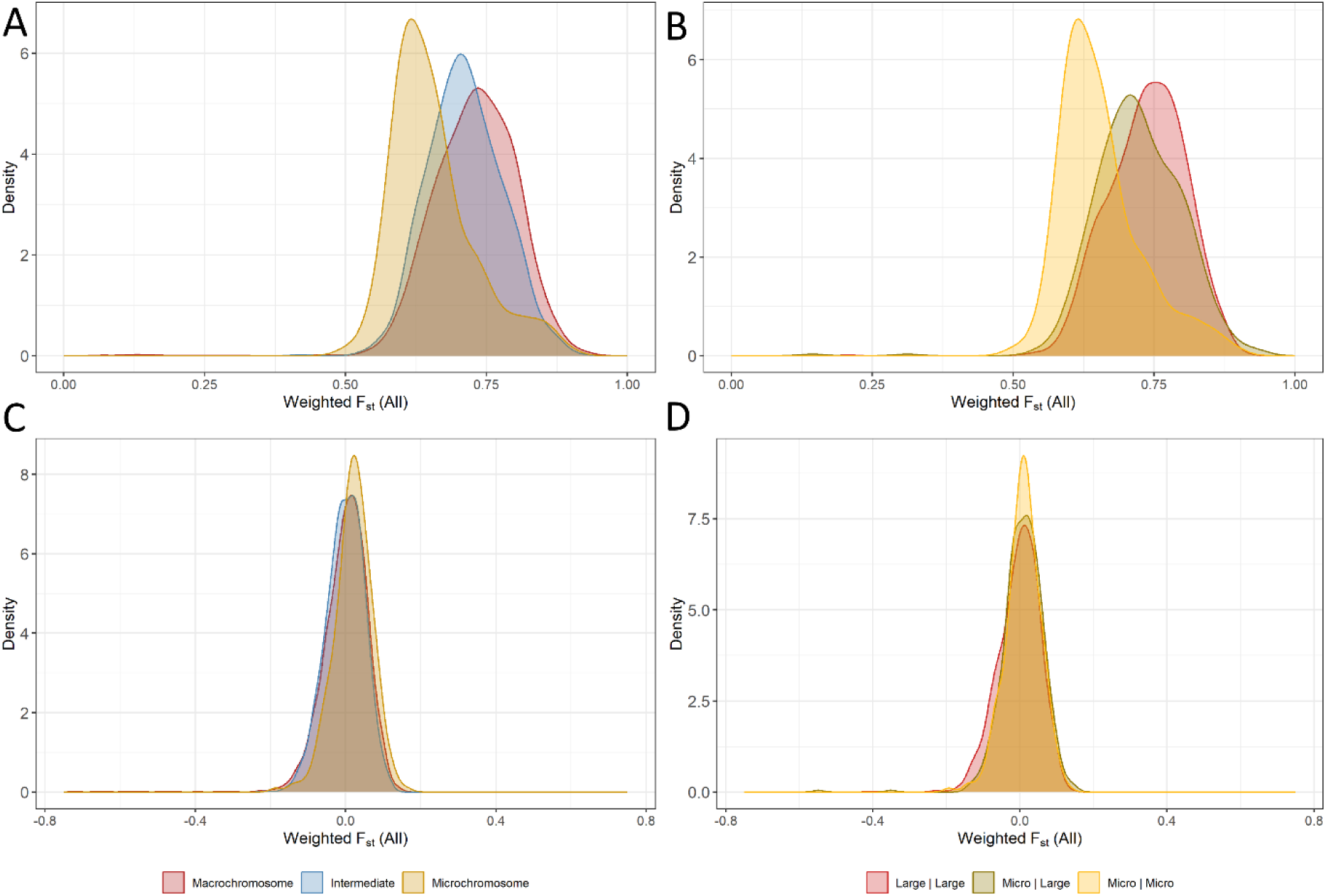
All falcon F_st_ by current and ancestral chromosome type. A) All falcon F_st_ by chromosome type; B) All falcon F_st_ by Ancestral | Current chromosome type. C) All falcon F_st_ by chromosome type after controlling for recombination rate. E) All falcon F_st_ by Ancestral | Current chromosome type after controlling for recombination.

Autozygosity may be linked to patterns of recombination and past demography. Autozygosity did differ by current chromosome state across all falcons, albeit with a low effect size (Kruskal-Wallace: χ^2^= 38.359; DF=2; p<10^-6^; ε²= 0.00418; R^2^= 0.00445. Pairwise Mann-Whitney U tests: p<10^-6^ for intermediate versus macrochromosomes). These results were driven by elevated levels of autozygosity on intermediate chromosomes relative to macrochromosomes in gyrfalcons (Pairwise Mann-Whitney U tests: p<10^-6^) and peregrine falcons (Pairwise Mann-Whitney U tests: p=0.000065). Peregrine falcons also displayed lower autozygosity on microchromosomes relative to intermediate chromosomes (p=0.0019999). In gyrfalcons regions conserved on microchromosomes had lower autozygosity than regions conserved on large chromosomes (p<10^-6^), but no other effects of ancestral state were observed. Autozygosity did not correlate to recombination rate, and controlling for recombination rate did not meaningful impact any autozygosity analyses.

Genomic differentiation was expected to be higher on large chromosomes due to reduced recombination. We assessed patterns of differentiation across falcon species using weighted Cockerham-Weir F_st_ (Weir & Cockerham, 1984). This method weights F_st_ by per locus contribution to total genetic variance, and was applied to all falcons and just the saker and gyrfalcons. We see that all chromosome types show significantly different distributions of F_st_ (Kruskal-Wallace: χ^2^= 313.89; DF=2; p<10^-6^; ε²= 0.03792; R^2^=0.03465. Pairwise Mann-Whitney U tests: p<10^-6^ for all), such that larger chromosome types tend toward greater differentiation than those smaller than them (Fig. 5C & D). The same pattern was found when ancestral state was considered (Kruskal-Wallace: χ^2^= 313.89; DF=2; p<10^-6^; ε²= 0.03792; R^2^=0.09581. Pairwise Mann-Whitney U tests: p<10^-6^ for Micro | Micro versus Large | Large and Micro | Large; p=0.0000083 for Micro | Large versus Large | Large), such that windows conserved on large chromosomes tend toward greater differentiation than those conserved on microchromosomes, with regions fused to large chromosomes from macrochromosomes tending toward intermediate values of differentiation to those conserved on large chromosomes and microchromosomes. As these findings are inversely parallel to patterns of recombination rate, we also ran these analyses on the residuals of the all-species F_st_ and recombination rate such that the lineage effects of recombination rate were regressed out. Recombination rate explained only a small fraction of differentiation (Linear Model: F=14.22; DF=8225; p<10^-6^; R^2^=0.003446) and results were effectively the same for current chromosome types when these residuals were used (Fig. 5E). However, the differences in differentiation between regions of microchromosomes fused to large chromosomes and regions conserved on microchromosomes appear to be primarily driven by recombination rate and no difference in F_st_ distributions detected between these after controlling effect of recombination rate. In contrast, these fusions displayed a reverse in sign relative to conserved regions on large chromosomes after controlling for recombination rate, such that microchromosomal fusions onto larger chromosomes now showed greater differentiation (Pairwise Mann-Whitney U tests: p=0.0002) than regions conserved on large chromosomes after controlling for recombination rate.

When we look at patterns of F_st_ between the saker and gyrfalcon we see similar but less pronounced results (Fig. 6). Clear differences in distributions of differentiation were found based on current chromosome state (Kruskal-Wallace: χ^2^= 122.8; DF=2; p<10^-6^; ε²= 0.014687; R^2^=0.01364. Pairwise Mann-Whitney U tests: p<10^-6^ for all macrochromosome comparisons; p=0.00069 for microchromosomes versus intermediate) with macrochromosomes and intermediate chromosomes tending toward higher levels of differentiation than microchromosomes (Fig. 6A). When ancestral state is considered (Kruskal-Wallace: χ^2^= 122.8; DF=2; p<10^-6^; ε²= 0.02036; R^2^=0.01226. Pairwise Mann-Whitney U tests: p<10^-6^ for Micro | Micro versus Large | Large; p=0.000084 for Micro | Micro versus Micro | Large; p=0.00016 for Micro | Large versus Large | Large), we see that regions conserved on large chromosomes tend toward higher levels of differentiation than those conserved on microchromosomes (Fig. 6B). Regions fused to large chromosomes from microchromosomes tend toward lower F_st_ than regions conserved on large chromosomes and higher F_st_ than regions conserved on microchromosomes. When we regress out the effect of recombination rate (Linear Model: F=1,660; DF=8,226; p<10^-6^; R^2^=0.168), we see that microchromosomes now show higher mean Fst (Pairwise Mann-Whitney U tests: p<10^-6^ for both) than either of the larger chromosomes types (Fig. 6C & D). Saker-Gyfalcon mean F_st_ differed significantly after controlling for recombination when ancestral state was considered (Kruskal-Wallace: χ^2^= 57.608; DF=2; p<10^-6^; ε²= 0.018934; R^2^=0.01233. Pairwise Mann-Whitney U tests: p<10^-6^ for both Micro | Micro comparisons), with regions conserved on microchromosomes showing higher levels of differentiation than those conserved on large chromosomes and fused to large chromosomes from microchromosomes.

**Figure 6:**
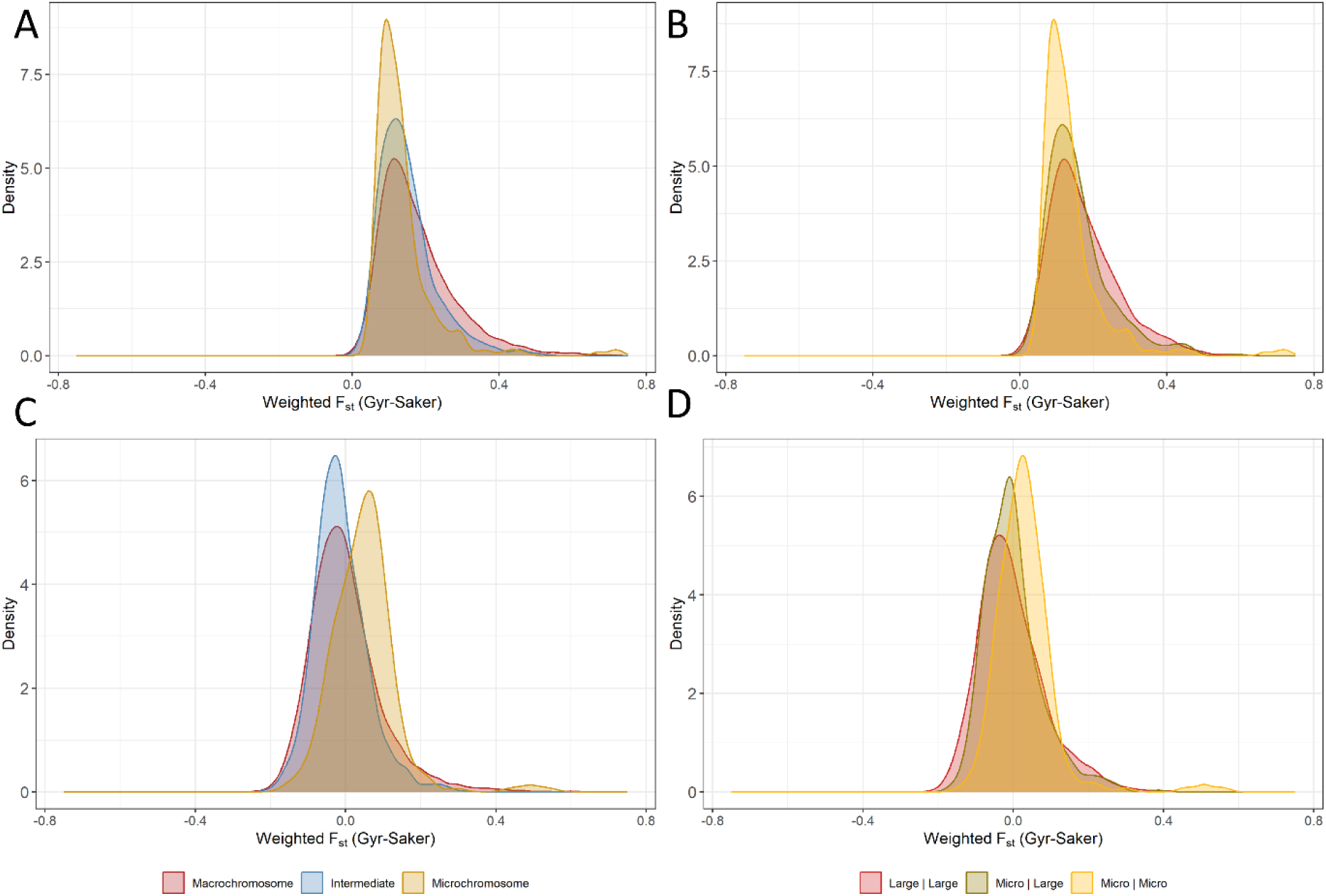
Saker-Gyrfalcon F_st_ by current and ancestral chromosome type. A) Saker-Gyrfalcon F_st_ by chromosome type; B) Saker-Gyrfalcon F_st_ by Ancestral | Current chromosome type. C) Saker-Gyrfalcon F_st_ by chromosome type after controlling for recombination rate. E) Saker-Gyrfalcon F_st_ by Ancestral | Current chromosome type after controlling for recombination.

Differences in genomic architecture and chromosomal fusions were suspected to have influenced patterns of selection. We assessed whether outliers for signals of selection were enriched on certain chromosome types using Fisher’s exact tests. There was no significant enrichment of outliers of selective sweeps or balancing selection on any current chromosome type relative to any other chromosome type, nor did any significant enrichments for indicators of selection occur based on ancestral state.

## Discussion

Here we analyzed the population genomics and patterns of divergence in large falcons in the context of genomic architecture. Our findings support previous reports suggesting relatively recent divergence of large falcons within the span of the last several hundred-thousand years (Fuchs et al., 2015; Wilcox et al., 2022; Zinevich et al., 2023). Though we cannot rule out any natural hybridization, our results support the distinctness and effective reproductive isolation of gyrfalcons, saker falcons, and peregrine falcons as species despite this recent divergence. We perform scans for selective sweeps and balancing selection in each of these large falcons and find unique candidate genes with patterns of diversity that are indicative of these selection regimes. We evaluate patterns of divergence in the context of genomic architecture, including chromosome types, and fusions of chromosomes in falcons. We confirm the predicted (Ellegren, 2010; Wilcox et al., 2022) correlation between recombination rate and chromosome types in birds, and show that fusions of chromosomes in falcon have lowered recombination rate in these regions. We find that current chromosome state and historical fusions influence almost all aspects of divergence between the large falcon species. These patterns are driven in part, but not entirely, by recombination rate. Taken together our findings reveal large falcons to be in the earliest stages of pre-zygotic isolation, and the indispensable importance of genomic architecture in population genomics (Campbell et al., 2018) of birds (Manthey et al., 2021; Manthey et al., 2016). Our findings do not, however, reveal a clear role for genomic rearrangement in falcon adaptation or speciation.

The three large falcons studied have diverged within the last several hundred thousand years and all show signs near-total reproductive isolation at one time—although interpretation of this result is hindered by our lack of knowledge of falcon origins. Admixture analyses indicate distinct genetic clusters corresponding to each species. Within-species multiple sequential Markov coalescents (from MSMC2) show distinct and consistent demographies within each species regardless of individuals sampled. Cross-species multiple sequential Markov coalescents likewise inferred a reduction in gene flow between all species over time. Gene flow appears to have continued between gyrfalcons and saker falcons until around, or just before, the beginning of the Holocene, approximately 12,000 years ago. Thus, the slow reduction in gene flow is consistent with contemporary tendencies toward parapatry in *Hierofalco* (Wink et al., 2004), and may be of interest to ongoing debates about the origins of the Altai falcon (Zinevich et al., 2023). Though our results do not preclude occasional natural hybridization between species (Moseikin & Ellis, 2004), they do suggest that this does not make any meaningful contribution to the population genomics of either the saker falcon or gyrfalcon species. Our results do, rather surprisingly, suggest introgression between peregrine falcons and the two *Hierofalco* species and/or their common ancestor. Given the weakness of MSMC2 at recent time scales, more sophisticated and focused techniques may be necessary to resolve the source, timing and impact of these introgressions. Further analyses of these introgression may be particularly interesting from the perspective of conservation and anthropogenic influences, given the major effects of captive-breeding and release on wild peregrine population (Wilcox et al., 2019) and their capacity to influence cross species hybridization (Lindberg & Nesje, 2002). They may also be interesting from the perspective of local adaptation and landscape genomics, given the extremely broad distribution of peregrine falcons and their wide variety of purported sub-species (Johnson et al., 2023; Wink, 2018). As such, deeper studies on peregrine-*Hierofalco* introgression would benefit from geographical information, which are lacking in the present study.

Falcons present an excellent system for study of speciation dynamics and their cultural and commercial importance making questions of falcon inter-relatedness important to legal practices, conservation, and the traditions of falconry. Our findings on all the large falcon species suggest that they are in earliest stages of pre-zygotic isolation. Previous estimates of divergence between peregrine falcons and *Hierofalco* have ranged from about 600,000 (Wilcox et al., 2022) to 2-3 million (Fuchs et al., 2015; Zhan et al., 2013). Previous estimates of saker falcon and gyrfalcon divergence have ranged from 34 (Pan et al., 2017) to 109 (Al-Ajli et al., 2023) to 149 (Wilcox et al., 2022) to 300 (Hu et al., 2022) thousand years ago. Our coalescent approach suggests earlier divergence times for the peregrine falcon from *Hierofalco*, but ambiguity in estimating saker and gyrfalcon divergence. Indeed, attempts at clear coalescent times are a flaw of phylogenetic approaches. Our data suggest that these species diverged in the presence of reducing gene flow, rather than through a sudden and strict imposition of allopatry. Birds are generally expected to speciate in allopatry and then reinforce following secondary contact (Winker, 2021). The extent and causes of geographic isolation necessary for avian speciation remain poorly resolved (Cooney et al., 2017; Winker, 2021), however, and these findings join a growing body of evidence (Barrera-Guzmán et al., 2022; Battey & Klicka, 2017; Beckman et al., 2018; Brown et al., 2025; Carling et al., 2010; Everson et al., 2019; Funk et al., 2021; Li et al., 2022; Linck et al., 2020; McLaughlin et al., 2020; Zarza et al., 2016) for speciation of birds in the presence of limited gene flow. Peregrines in contrast seem to have reestablished minimal gene flow with *Hierofalco* following some isolation, another increasingly well documented phenomenon in birds (Brown et al., 2025; Li et al., 2022; Musher et al., 2022; Weir et al., 2024). The recent divergence of the saker and gyrfalcon and the permeability of peregrine falcons to introgression with *Hierofalco* demonstrate the utility of falcons for studies on speciation. Ongoing studies on the biogeography of falcon speciation may contribute to an improved understanding of falcon and avian speciation in general. Regardless, our results demonstrate that peregrine, saker, and gyrfalcons are distinct as species and can be delineated using standard population genomic techniques. This has clear implications to traditional and modern applications of falconry and the international falcon trade (Wilcox et al., 2019).

All falcon species are sufficiently diverged to show unique patterns of selective sweeps and balancing selection. Though clear morphological and ecological differences have been identified between these falcons (Cade, 2011; Wilcox et al., 2019), our findings suggest that recent and strong directional evolution in large falcons has occurred primarily at the level of cellular process and mechanisms. Scans for selective sweeps are relatively crude (Soni et al., 2023; Szpak et al., 2019), and it is unclear how these differences may scale up to observed phenotypic differences. Large falcons, due to their very recent divergence, blur the line (Yeaman, 2022) between local adaptation (Tigano & Friesen, 2016; Yeaman, 2015) and speciation (Stankowski & Ravinet, 2021). Finer studies using environmental variables, morphometrics, and techniques focused on polygenic adaptation may be better suited to identifying divergent selective pressures on these large falcons (Barghi et al., 2020; Stephan, 2016). The small number of genes under balancing selection, affirm previous hypotheses of the general rarity of balancing selection (Hedrick, 2012), although methodological advances may allow detection of additional examples (Bitarello et al., 2023; Bitarello et al., 2018). These genes all seem to relate to metabolic, developmental, or physiological tradeoffs. We notably do not find MHC to be under balancing selection, in accordance with previous studies on falcons (Gangoso et al., 2012); nor do we see any clear enrichment of immune genes, as suggested in primates (Bitarello et al., 2023; Bitarello et al., 2018). E3 ubiquitin-protein ligase SMURF1 has pleiotropic effects (Wan et al., 2023) and could relate to trade-offs in the realms of immunity, development, or both. Aspartate aminotransferase has dual effects on amino-acid metabolism and respiration (Ndrepepa, 2021; Wang & Xu, 2025) and could experience trade-offs between these. Its expression levels have been reported to differ between the large falcon species used in this study and their hybrids (Lierz & Hafez, 2006). Cytoplasmic actin has a very wide range of functions (Pollard, 2016; Svitkina, 2018) including, cell development, mechanical performance, immune functions, and even gene expression (Percipalle & Vartiainen, 2019). It has been reported to be under diversifying selection—expansions and specialization and/or divergent selection between lineages—in algae (Wu et al., 2009), plants (Slajcherová et al., 2012), amoeba (Lahr et al., 2011), and primates (Zhu et al., 2013), although not in birds. Its ubiquitous role in a multitude of phenotypes make its regime of balancing selection plausible but mysterious. Regardless, scans for selection highlight shared evolutionary pressures within falcons and a potential role for housekeeping genes during their adaption.

The primary focus of this work is on the role of genomic architecture in adaption and speciation. We do not find any specific role for chromosome type or chromosomal fusions in adaption: Genes under selective sweeps and balancing selection are not enriched on any particular chromosome type. The influence of directional and balancing selection on any given genes can presumably be highly variable over evolutionary time (Bitarello et al., 2023; Soni et al., 2023; Szpak et al., 2019), and the capacity for genomic architecture to favor any selective regime may be too nuanced for detection by our approach. We nonetheless find strong associations between all population genomic indicators and chromosome type. Recombination rate and nucleotide diversity are higher on smaller chromosomes. Despite an overall correlation across species, higher recombination rates are not necessarily predictors of higher π, and indeed macrochromosomes do not necessarily have lower levels of diversity than intermediate chromosomes. Variation in diversity (π) and recombination rate were also much higher on microchromosomes than other chromosome types, although this trend was much more strongly pronounced for recombination rates than for π. Though these general patterns reflect those observed previously in other birds from a comparative genomic perspective (Axelsson et al., 2005; Ellegren, 2010), our results provide an assessment of these features from a population genomic perspective. They additionally provide novel insights into the roles of chromosomal fusions in altering population genomic signatures.

Falcons provide a system to assess how chromosomal fusions impact population genomics and species divergence (Wilcox et al., 2022; Wilcox et al., 2019). We find that fusions of smaller chromosomes to larger chromosomes lower recombination rate, but only to an intermediate level—fused regions of microchromosomes do not converge to the recombination rate of large chromosomes, presumably due to either the regions of chromosomes to which they fuse or some sequence variation on these regions. As such these findings are consistent with prior suggestions of sequence-based hotspots of recombination in birds (Kawakami et al., 2014; Singhal et al., 2015; Tiemann-Boege et al., 2017; Topaloudis et al., 2024). The observed reductions in recombination rate can, nonetheless, explain our previous observations of decreasing GC content and CpG sites in these fused regions (Wilcox et al., 2022) through reduced conversion (Mugal et al., 2015). This provides for the possibility that chromosomal fusions may have also had a major impact on epigenetic regulation between falcons as a result of CpG hypermutability (Francioli et al., 2015; Nesta et al., 2021) and reduced recombination in these regions (Paudel et al., 2020), although this hypothesis lies beyond the scope of our current study. Fusions from microchromosomes do, however, result in regions of increased diversity (π) on macrochromosomes and intermediate between large chromosomes and microchromosomes. This suggests altered substitution regimes (Kimura, 1968) or sequence-related patterns of substitution (Barroso & Dutheil, 2023) that are retained following fusion. Overall, our results are similar to those reported recently in butterflies, which are attributed to changes in background selection regimes (Cicconardi et al., 2021; Nasvall et al., 2023). If fused regions were under reduced selective constraints, then background selection should be reduced as well. We nonetheless show that these effects apply to other indicators as well. Fusions also result in decreased Tajima’s D in all falcons and increased F_st_ between the recently diverged saker and gyrfalcons. This indicates a capacity for past chromosomal rearrangements to influence contemporary assessments of regions under selection. It also provides for a role of genomic architecture in species divergence.

Our research describes a pattern of divergence in falcons that is heterogeneous across the genome and chromosomal context. Microchromosomes trend toward higher diversity and disproportionately contribute to within species diversity. Lower recombination on intermediate and macrochromosomes enhances drift and accelerates lineage sorting. Fusions further alter patterns of divergence by creating regions of accelerated divergence with altered recombination, substitution, and fixation regimes. Our study deals primarily with patterns of variation and lineage sorting. Molecular mechanisms of genomic incompatibilities are unknown in falcons—and as all three species in this study produce viable offspring in captivity (Lindberg & Nesje, 2002; Wilcox et al., 2019)—almost certainly arise more from mate choice and selective regimes than negative epistasis at the molecular level. Regions of genomic incompatibility are now being better identified in birds and other natural populations (Anderson & Weir, 2022). Improved annotation of such regions in falcons—for example, through analysis of hybrids—may help to better elucidate the true impacts of chromosomal fusions on falcon speciation and evolution.

Regardless, our study links population genomics and divergence to genomic architecture in large falcons. We find that these are inseparable, and that simply controlling for recombination rate is not sufficient to explain variation in population genomic indicators and genomic heterogeneities. Our results join with emerging studies on the role of genomic architecture in avian speciation (Manthey et al., 2021; Manthey et al., 2016). Falcons are unusual among birds for their level of chromosome loss (Ellegren, 2010; O’Connor, Romanov, et al., 2018; Wilcox et al., 2019), and the extent to which past chromosome state influences population genomic patterns in other birds remains to be seen. Intrachromosomal rearrangements are, however, known to be much more common than previously anticipated in birds (Griffin et al., 2025; Huang et al., 2022; Kawakami et al., 2014; O’Connor, Farre, et al., 2018; O’Connor et al., 2024), and chromosomal rearrangements are relatively common in many other amniotes (Damas et al., 2021; Ellegren, 2010; Mezzasalma et al., 2024). As such, our results may provide some insight into the need to control for genomic architecture in other systems. The roles of genomic architecture on evolution within and between populations and species are of increasing interest (Campbell et al., 2018; Lucek et al., 2022; Manthey et al., 2021; Manthey et al., 2016). Large falcons provide an excellent system for assessing genomic architecture and speciation. Stronger mechanistic inference will require more representative sampling in the future.

## Methods

### Library Preparation and Sequencing

Blood samples were obtained from 14 gyrfalcons, 10 saker falcons, and 11 peregrine falcons by practicing veterinarians at Al Aseefa falcon hospital in Dubai, United Arab Emirates, during the course of normal veterinary care. The geographic origin of falcons in this study is unknown. Samples were extracted using the Qiagen MagAttract Extraction Kit (Hilden, Germany) following standard tissue extraction procedures except for a modification to the tissue lysis stage such that 25µl of blood was incubated in 175µl of ATL with 20µl of Proteinase K for 60 minutes at 56°C. Library preparation was performed with the 10X Chromium Genome High Throughput (HT) Gel and Bead Kit Version 2 (Pleasanton, CA, USA), barcoded with the Chromium i7 Multiplex kit, and sequenced on an Illumina Novaseq6000 using the NovaSeq6000 S4 300 Cycle Reagent Kit (San Diego, California, USA).

### Read Processing and Alignment

Linked-reads were aligned to our previously assembled (Wilcox et al., 2022) scaffold-scale lanner falcon reference genome (GCA_024433245.1) using the 10X Genomics Longranger software (Marks et al., 2019). The lanner falcon was chosen as it provided the best opportunity for a recent outgroup sequence to all included falcon species in the absence of high-quality assemblies from hobbies or the prairie falcon (Fuchs et al., 2015) — Longranger does not perform well with more distant reference genomes such as those from available kestrel assemblies. Variant calling was performed with GATK (version 3.5) in accordance with the recommendations of the 10X Genomics Longranger pipeline. Variants were filtered to remove those with overall Phred scores <30, Phred scores <15 for any heterozygous allele, Phred scores <50 for any homozygous allele, an allele fraction of <15%, inconsistent phasing, >3bp of unphased homopolymer insertions, and those that aligned poorly to the surrounding sequence or to duplicated loci using linked-read information. Unless otherwise noted, variants were further filtered to remove those at CpG sites in any conspecific sequence. Finally, all scaffolds less than 100KB (including the mitochondrial scaffold) and scaffolds potentially belonging to sex chromosomes were removed from all analyses. Scaffolds were considered to potentially belong to sex chromosomes if their best alignment (based on most aligned BP) to the archival NCBI chromosome-scale gyrfalcon assembly “bFalRus1.pri” (GCF_015220075.1) occurred on the annotated Z or W chromosome. Several low coverage somatic mutations were initially detected in all species, but *all* of these were found at CpG sites and removed by the above filtration criteria.

### Summary Statistics

Summary statistics were computed on filtered genome alignments for each species. Fixed sites within species were excluded from analyses. Allele frequency spectra were computed in Plink 2.0 version v2.00a2.3LM (Chang et al., 2015). Nucleotide diversity (π) and Tajima’s D were computed on 100,000 bp windows using VCFtools version 0.1.14 (Danecek et al., 2011). Runs of homozygosity were calculated in BCFtools version 1.3.1 (Narasimhan et al., 2016).

Recombination rate was calculated using LDhat version 2.2 (Auton & McVean, 2007). Likelihoods tables were calculated for each species and parametrized using Theta estimates based on the published zebra finch mutation rate of 4.9 x 10^-9^ substitutions per base per generation (Smeds et al., 2016) and previously published effective population size estimates of each species obtained with PSMC (Wilcox et al., 2022). Phased haplotype sequences were produced for each scaffold as input using the “consensus” option in BCFtools. Recombination rates were estimated using the RhoMap model, which incorporates variation in recombination rate across scaffolds with estimated hotspots of recombination on each scaffold. RhoMap is a Bayesian process and was run on each scaffold separately with 4,200,000 iterations and sampled at intervals of 4,000 iterations following a burn-in of 1,200,000 iterations.

### Admixture and Demography

Admixture and demographic analyses were conducted within and across species as appropriate. PCA was conducted across all species in Plink 2.0. An admixture analysis was conducted across all species using Admixture version 1.3.0 (Alexander & Lange, 2011). The distributions of variants private and shared across species were calculated with the BCFtools “intersect” command. Demographic analyses within species were performed using MSMC2 version 2.1.1 (Schiffels & Wang, 2020). As MSMC2 is too computationally intensive to run on all samples at once, four phased haplotypes were subsampled at random from each species and one hundred runs were performed on 100 bootstraps of four haplotypes for each species to utilize all data and obtain an estimate of error arising from sample choice. MSMC2 was run both within species and with cross-species comparisons to assess for interbreeding. The ratio of ρ to θ was estimated to be 0.3 based on an estimated per generation median recombination rate in zebra finch of 1.9 x 10^-9^ recombination events per base pair (Singhal et al., 2015) per generation and an estimated per generation mutation rate of 4.9 x 10^-9^ mutations per base pair per generation in the collared flycatcher (Smeds et al., 2016).

### Selection and Differentiation

Scans for selective sweeps were performed using Sweepfinder2 (1.0) with allele frequency spectra calculated in Plink2.0 (see *Summary Statistics*) and data treated as non-polarized. Tests sites were performed at approximately 10,000 windows (“–lrg 10000”). The analysis controlled for recombination using a recombination map created from our above LDhat RhoMap and species-specific estimates of effective population size from our MSMC2 analysis (see *Admixture and Demography*). BetaScan was run using folded site frequency spectra. Tests were performed on 100,000 base pair windows with θ inferred as above. Divergence times were parametrized using previously published genome-wide estimates phylogenetic estimates (Wilcox et al., 2022) and effective population sizes were calculated using MSMC2 as above. F_st_ (Weir-Cockerham) was computed for 100KB windows using VCFtools (Danecek et al., 2011). Cutoff values for both α for selective sweeps (from SweepFinder2) and Beta for balancing selection (from BetaScan2) were z-transformed and deviations with z-scores greater than 5.2 were taken as candidates. Under a normal distribution this would correspond to a p-value of 10^-7^.

### Genomic Architecture

Genomic positions on current and former macrochromosomes, intermediate chromosomes and microchromosomes were annotated as described previously (Wilcox et al., 2022). In brief, current chromosome state was annotated by MUMmer alignment to archival chromosome-scale reference genomes for the gyrfalcon and peregrine falcon. Ancestral chromosome state was annotated by MUMmer alignment to chromosome-scale assemblies for the Swainson’s Thrush (RefSeq Accession: GCF_009819885.2), Kakapo (RefSeq Accession: GCF_004027225.2) and Seriema (NCBI Accession: GCA_009819825.1)—representing the two closest extant clades to Falconidae and their outgroup respectively. Annotations and analyses in the context of chromosome state were performed by 100kb windows such that a window needed to align consistently to a current or ancestral chromosome state to be included in respective analyses. Windows that aligned consistently to chromosome types within falcons but not consistently among relatives of Falconidae were removed from analyses involving ancestral state but not those involving current state.

### Statistics

Statistical analyses were performed in R sersion 3.5.1, 4.2.0, 4.4.3 and 4.5.2 using R studio (RStudio Team, 2022). An alpha value of 0.002 was used based on a Dunn-Šidák correction of a 0.05 alpha for 19 independent tests. Groups of pairwise tests used Bonferoni corrected p-values.

## Data Availability Statement

All data will be made available at time of publication.

## Acknowledgments

We would like to thank Dr. Peter McKinney of Al Aseefa Falcon Hospital for his support during the early stages of this project and for his helpful discussions. This work was supported by Tamkeen under the NYU Abu Dhabi Research Institute Center for Genomics and Systems Biology award (ADHPG-CGSB) to Stephane Boissinot and Youssef Idaghdour. We also acknowledge Marc Arnoux (NYUAD Sequencing Core) for his work on sequencing and Nizar Drou (NYUAD CGSB Bioinformatics Core) for providing bioinformatics assistance.

