## Supplementary Figures for "Genomic Architecture, Differentiation, and Adaptation in Three Large Falcons"

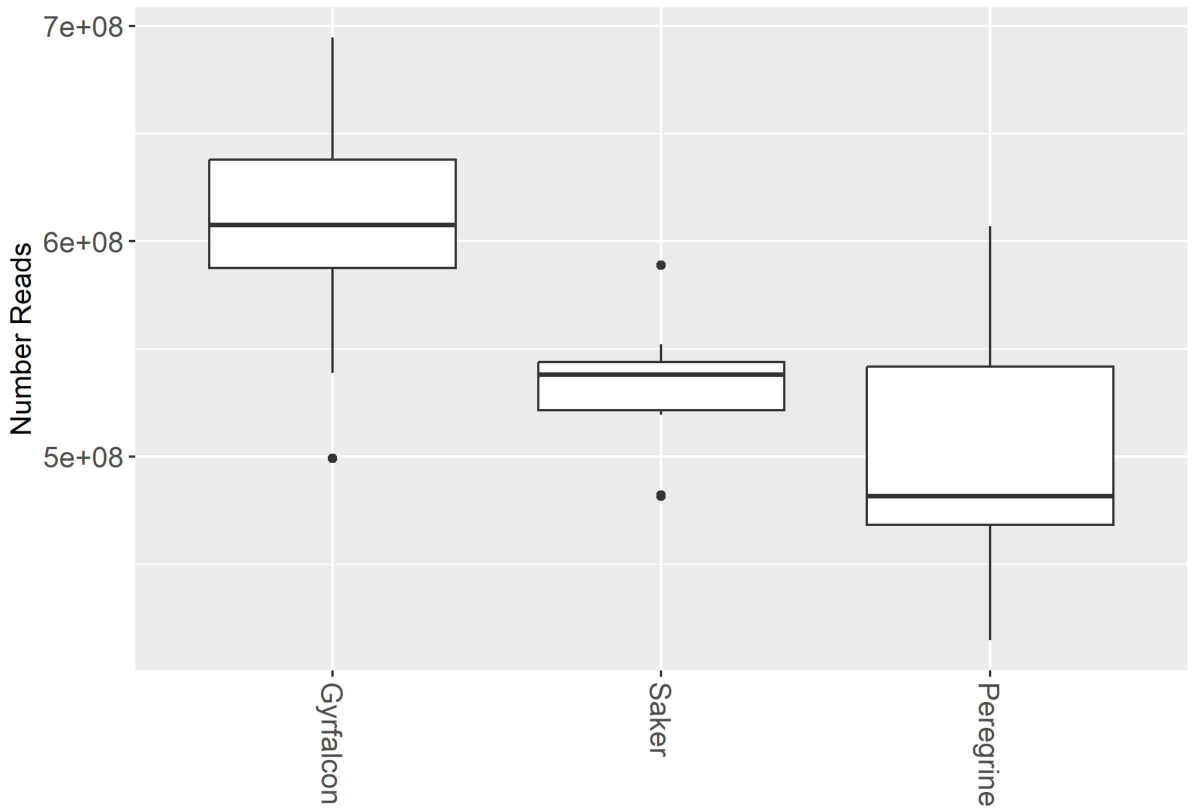


Supplementary Figure 1: Number of reads per sample by falcon species. Significant differences were observed between average read counts between species (Kruskall-Wallace Test: Kruskal–Wallis $\chi^2 = 16.769$; DF=2; p=0.0002283; $\varepsilon^2 = 0.46154$; $R^2$=0.4751) with gyrfalcons (Pairwise Mann-Whitney U tests: p=0.0019) having more reads than peregrine falcons.

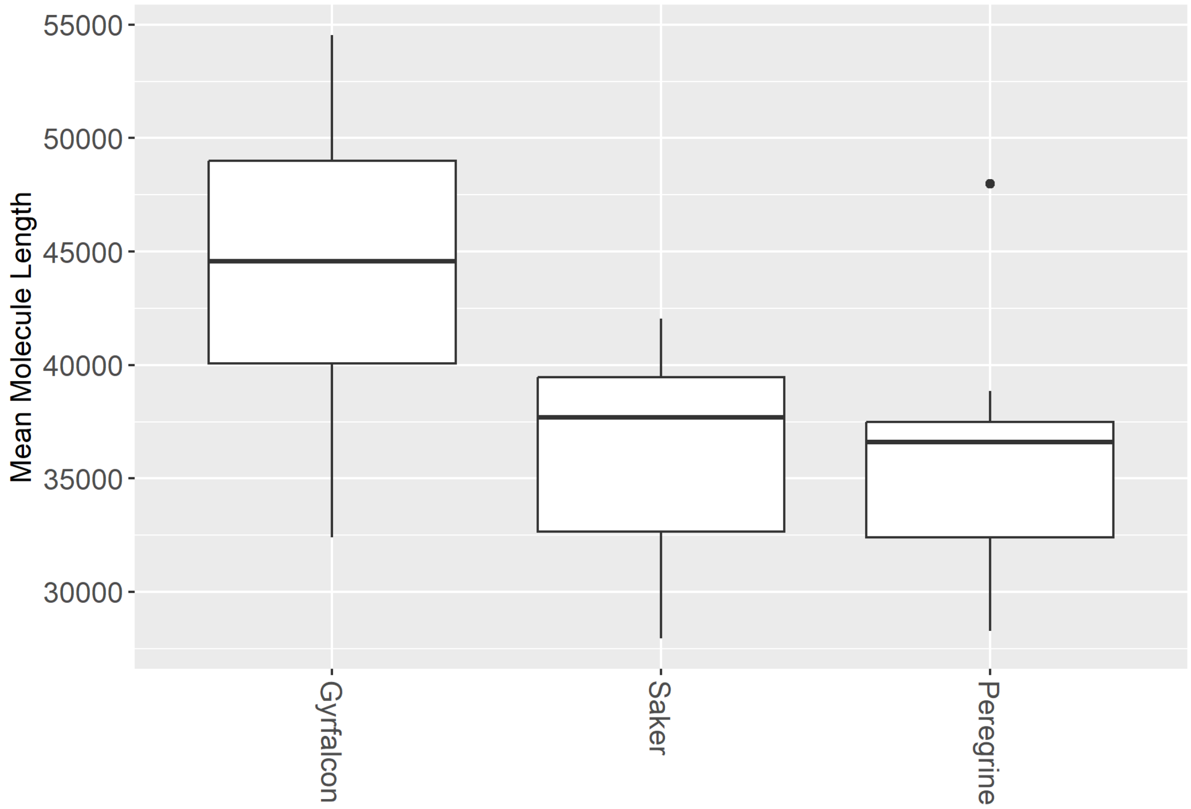


Supplementary Figure 2: linked-read molecule length by species. DNA molecule length did not differ significantly (Kruskall-Wallace Test: Kruskal–Wallis $\chi^2$ = 12.344; DF=2; p= 0.002087; $\varepsilon^2$=0.32324; $R^2$=0.3858).

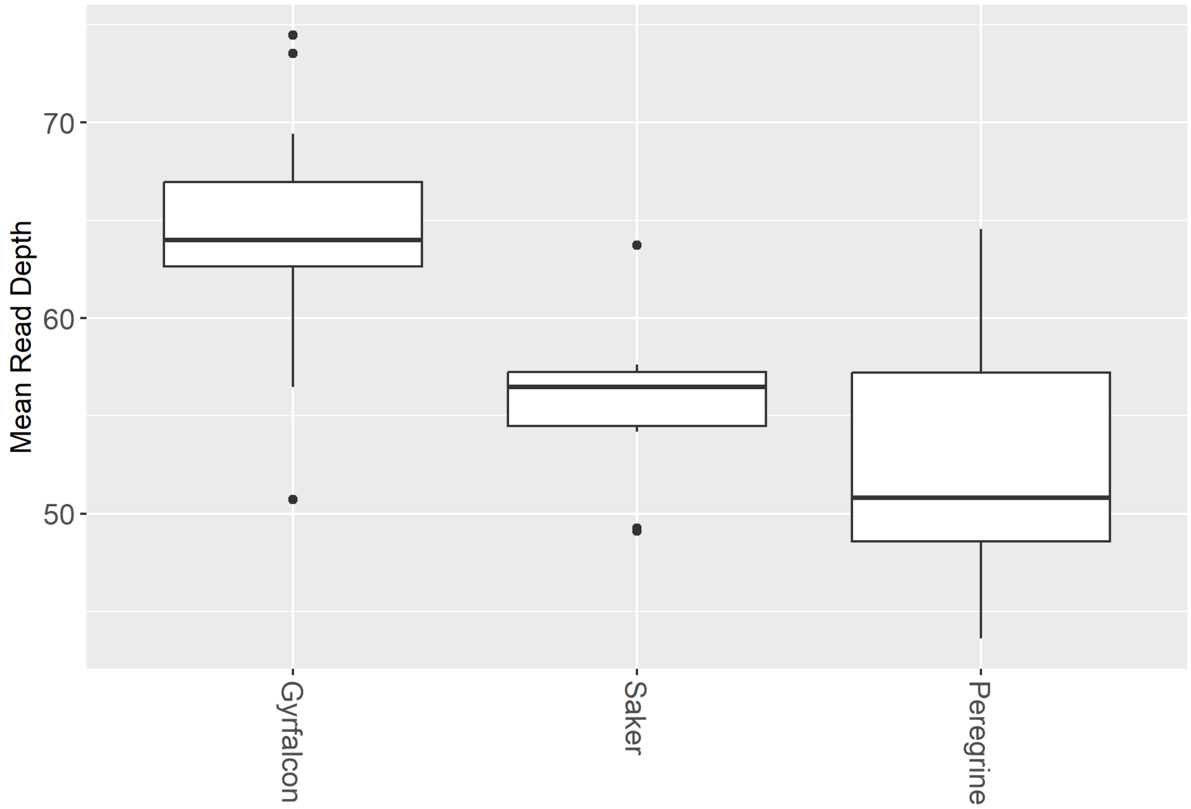


Supplementary Figure 3: mean aligned read depth per sample differed across species. (Kruskall-Wallace Test: Kruskal–Wallis $\chi^2$=15.415; DF=2; p=0.0004494; $\varepsilon^2$=0.41922; $R^2$=0.4307).

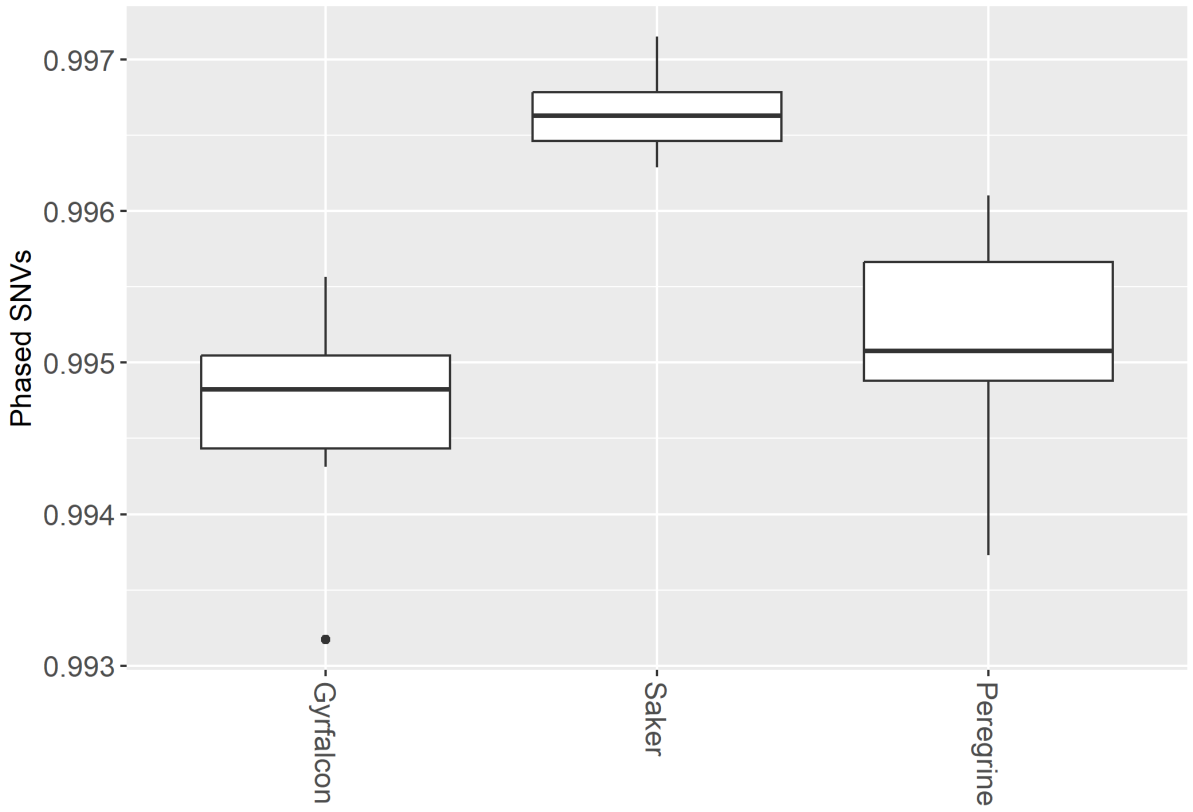


Supplementary Figure 4: Proportion of SNVs phased per sample by species. Significant differences were observed in phasing success across species (Kruskall-Wallace Test: Kruskal–Wallis $\chi^2$=22.734; DF=2; p=0.00001157; $\varepsilon^2$=0.64792; $R^2$=0.6949) with SNVs in the saker falcon phasing slightly better than those in the gyrfalcon (Pairwise Mann-Whitney U tests: p=0.00014) and peregrine falcon (Pairwise Mann-Whitney U tests: p=0.000017).

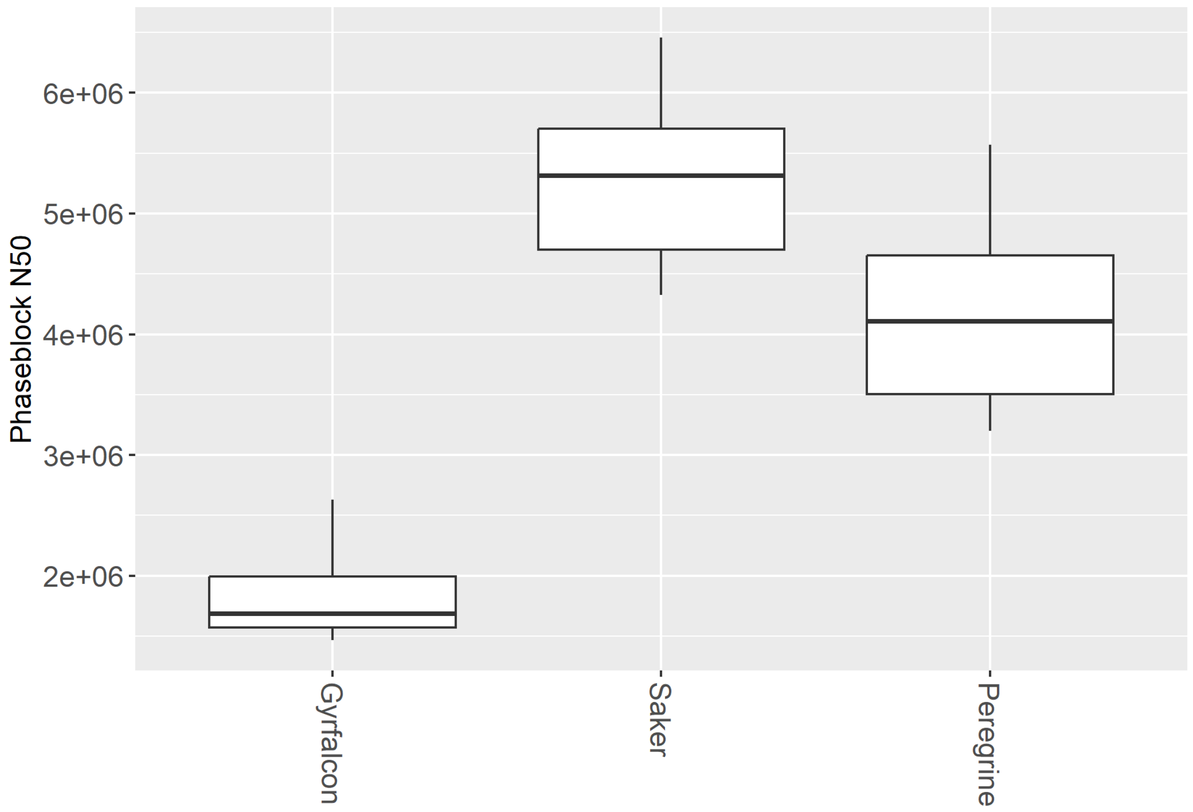


Supplementary Figure 5: Phase block N50 (in base pairs) across samples by species. Significant differences were observed between all species (Kruskall-Wallace Test: Kruskal–Wallis $\chi^2$= 27.417; DF=2; p=0.00000113; $\varepsilon^2$=0.79427; $R^2$=0.8683), with the saker (Pairwise Mann-Whitney U tests: p=0.00014) and peregrine (Pairwise Mann-Whitney U tests: p=0.0000084) falcons phasing better than gyrfalcons.

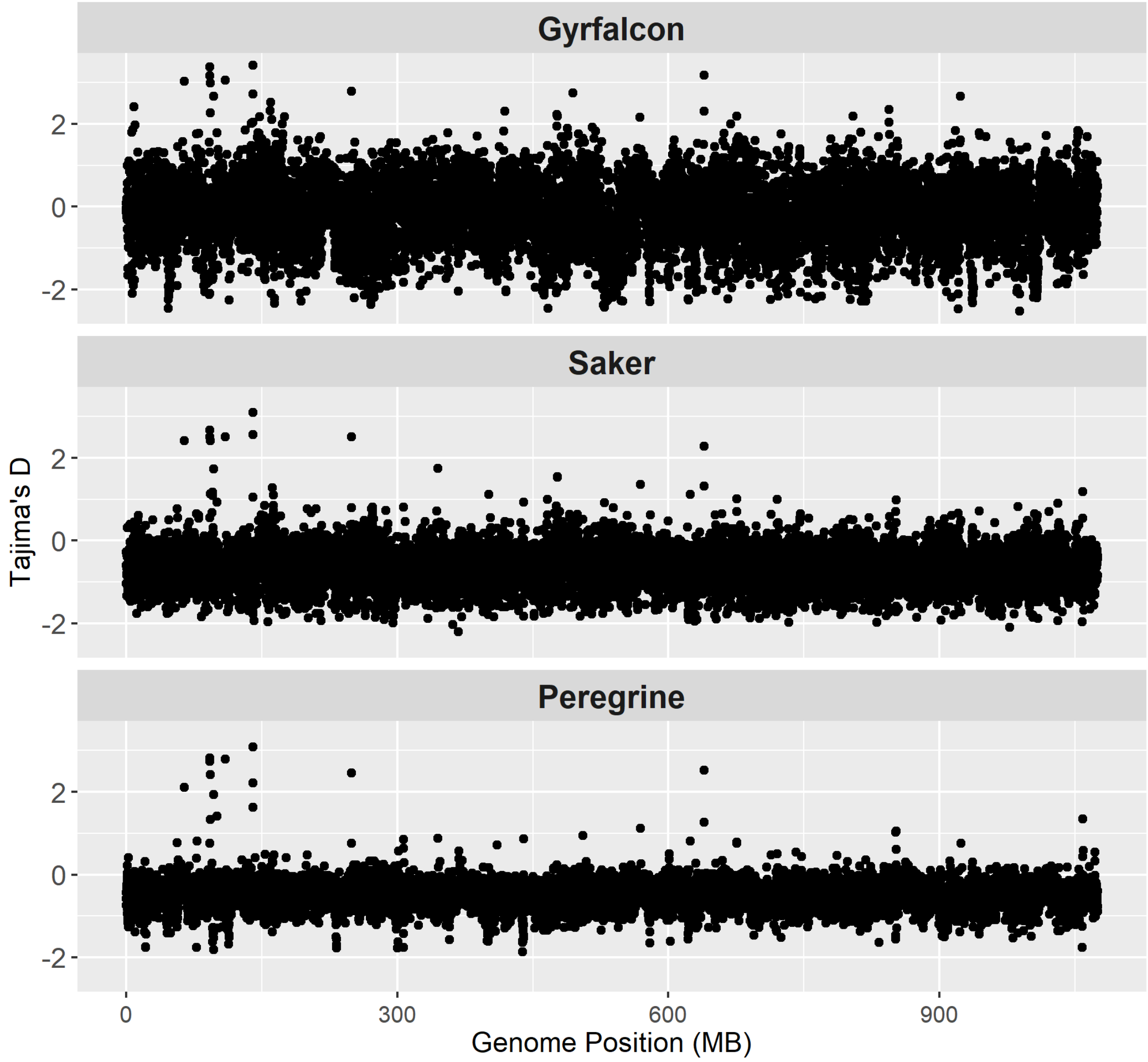

Supplementary Figure 6: Tajima's D of aligned 100KB windows faceted by species. All alignment plots in this manuscript share the same window positions.

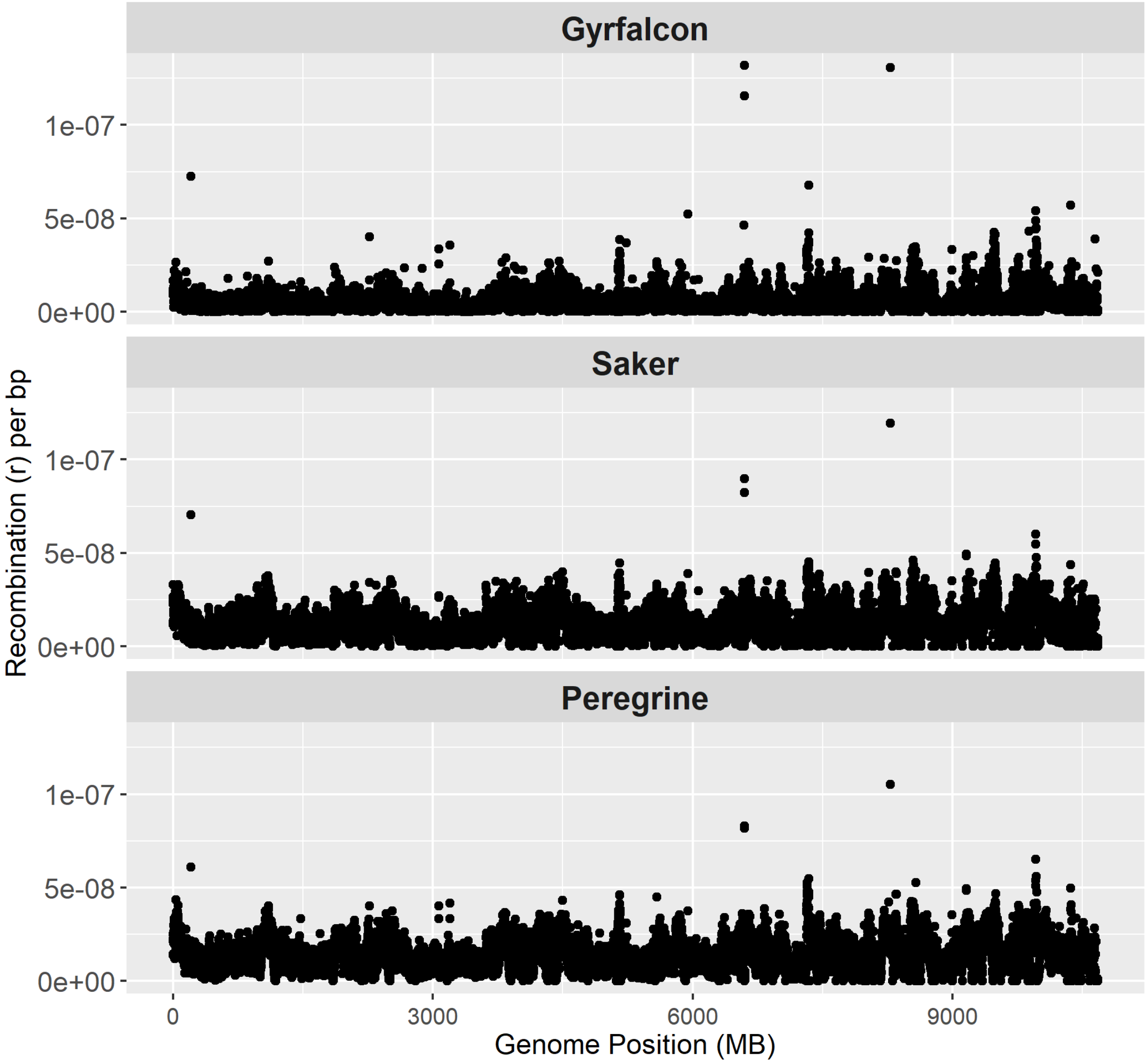

Supplementary Figure 7: Recombination rate, r, of aligned 100KB windows faceted by species. All alignment plots in this manuscript share the same window positions.

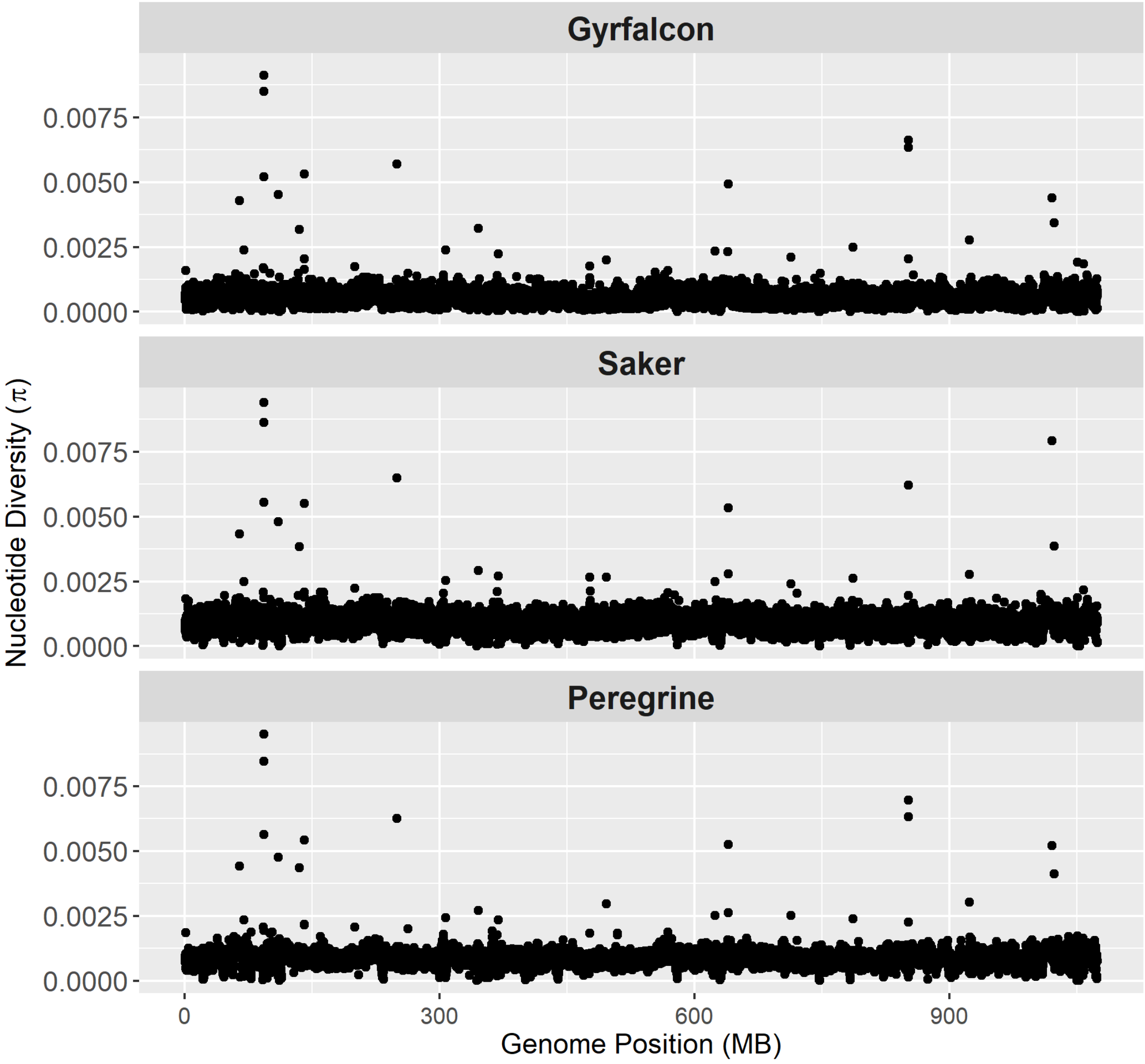


Supplementary Figure 8: Nucleotide diversity, π, of aligned 100KB windows faceted by species. All alignment plots in this manuscript share the same window positions.

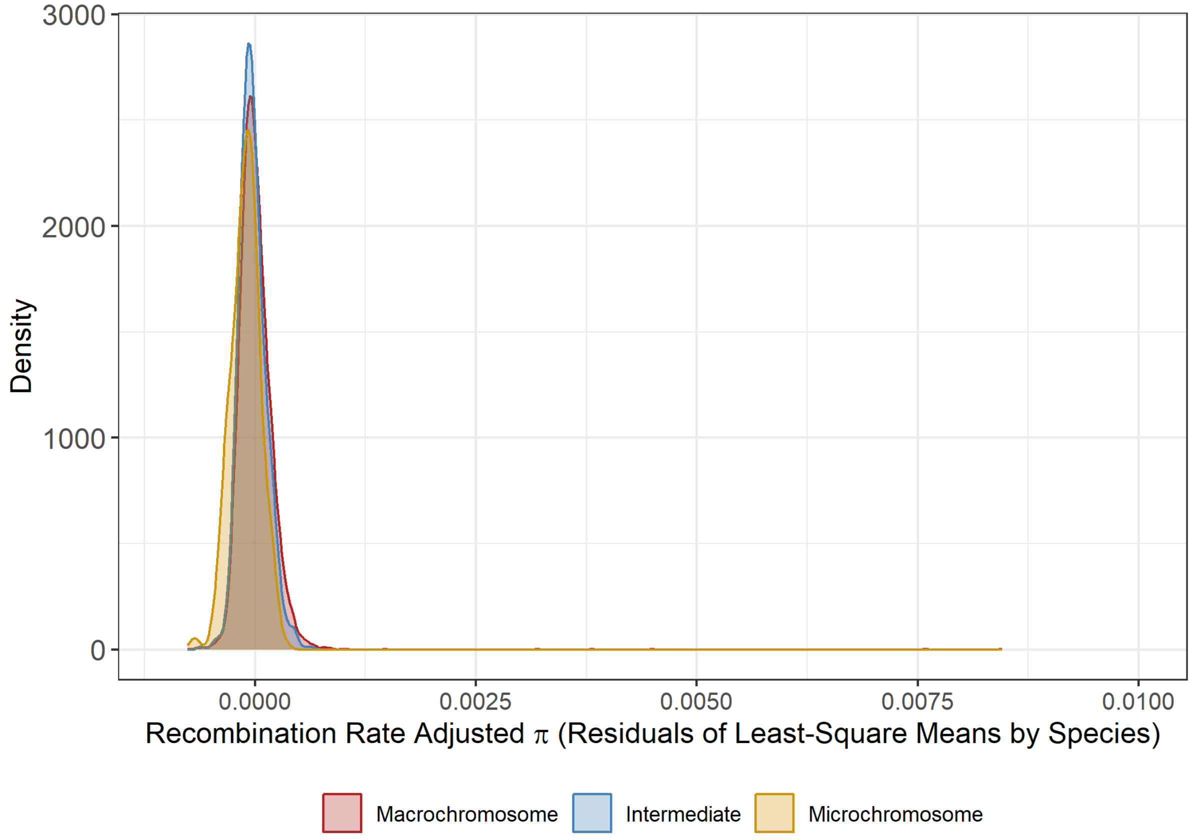


Supplementary Figure 9: Densities of residual nucleotide diversity, $\pi$, across 100KB windows by current chromosome type after regressing first on species (least square means) then on recombination rate.

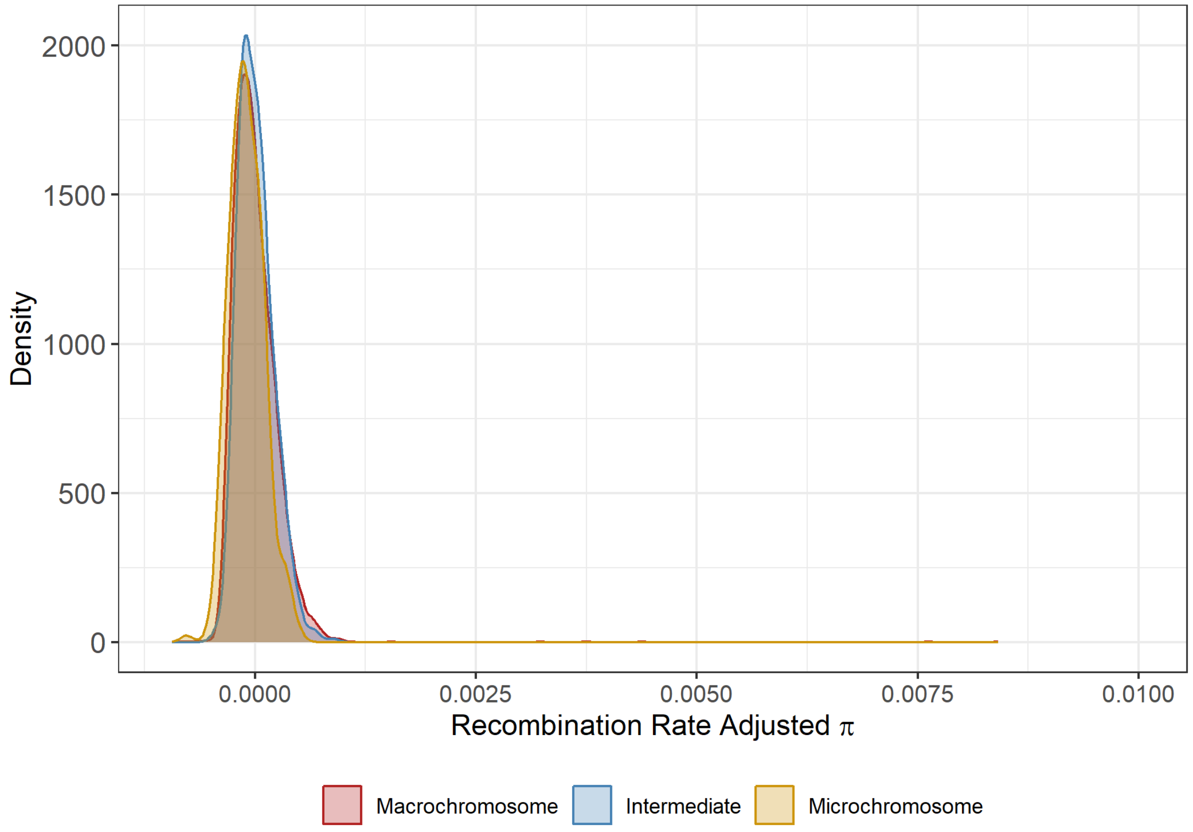


Supplementary Figure 10: Densities of residual nucleotide diversity, $\pi$, across 100KB windows by current chromosome type in gyrfalcons after regressing on recombination rate.

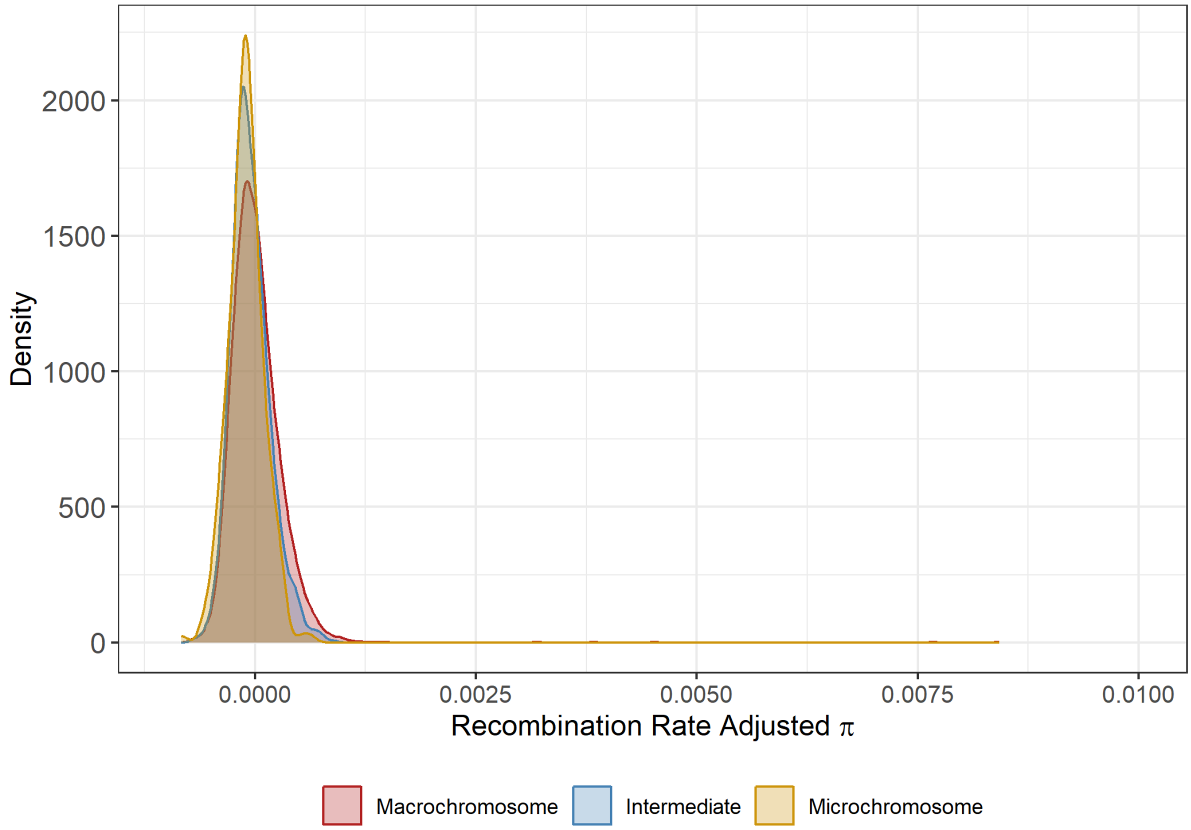


Supplementary Figure 11: Densities of residual nucleotide diversity, $\pi$, across 100KB windows by current chromosome type in saker falcons after regressing on recombination rate.

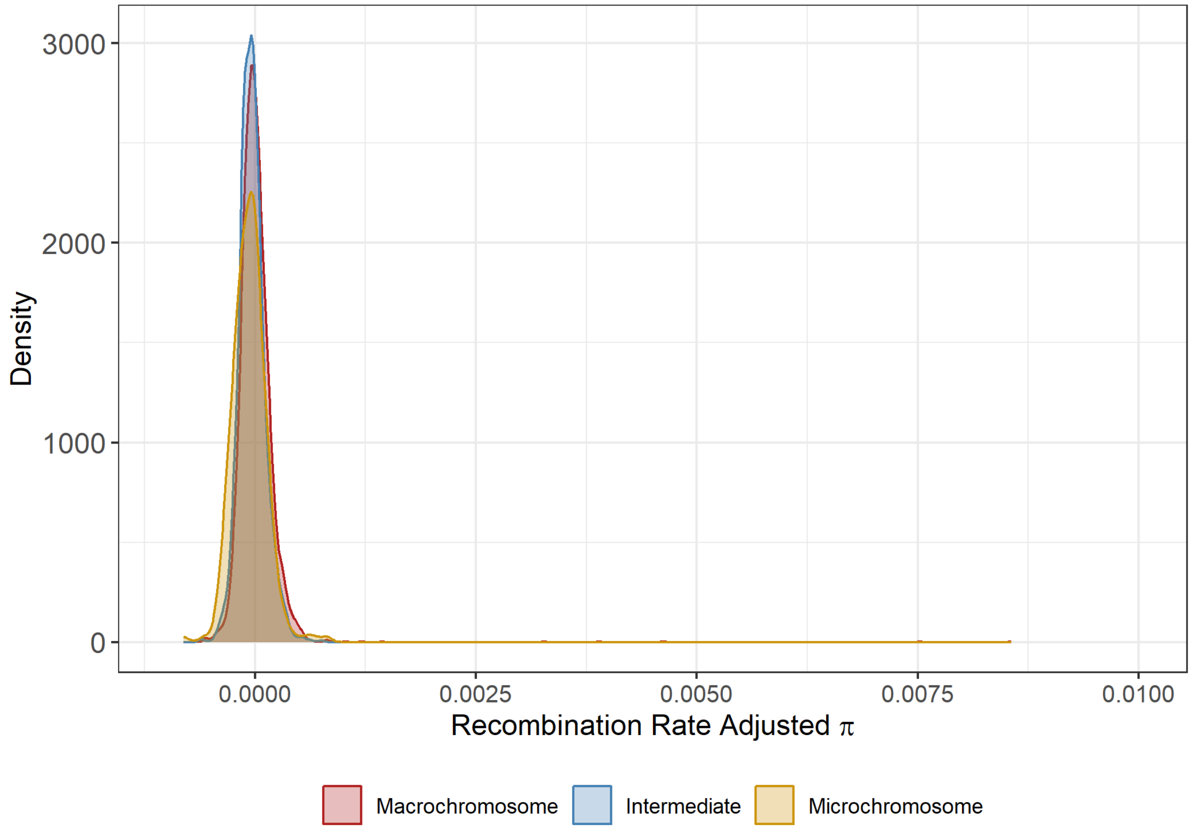


Supplementary Figure 12: Densities of residual nucleotide diversity, $\pi$, across 100KB windows by current chromosome type in peregrine falcons after regressing on recombination rate.

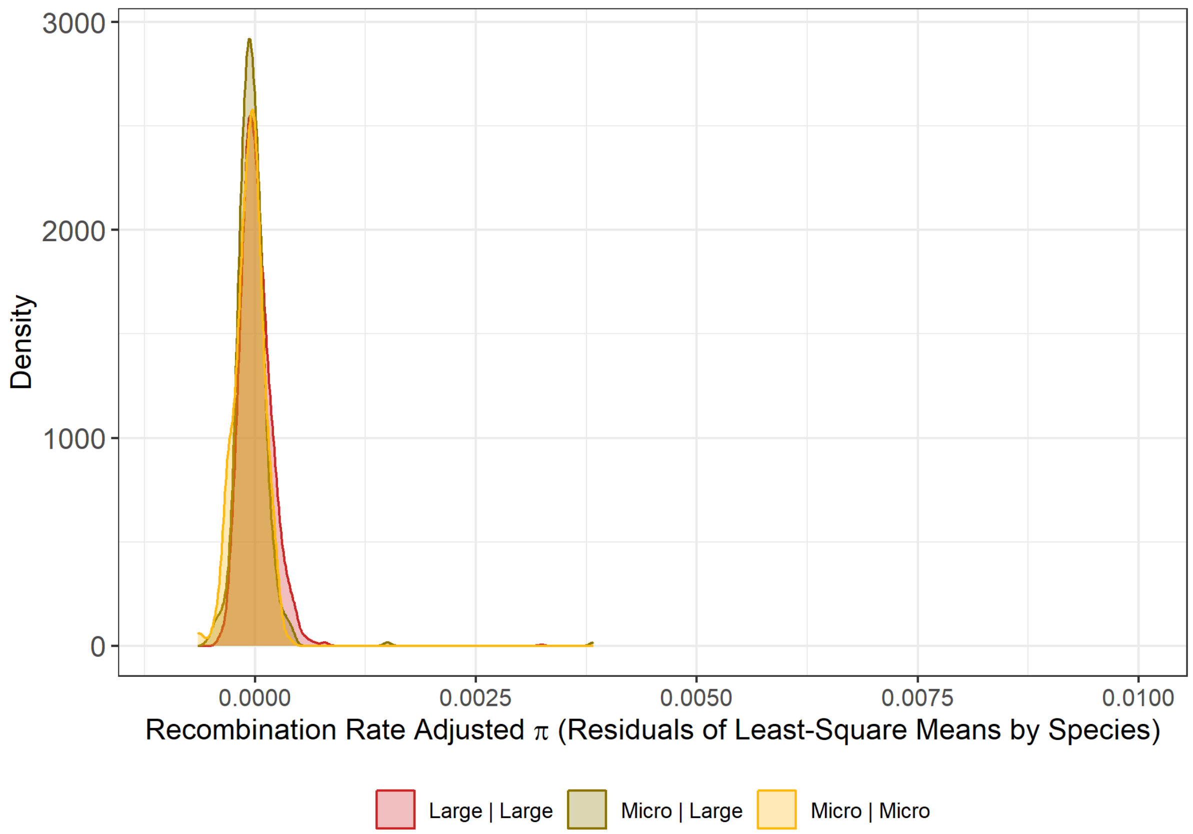


Supplementary Figure 13: Densities of residual nucleotide diversity, $\pi$, across 100KB windows by chromosomal history (Past | Current) after regressing first on species (least square means) then on recombination rate.

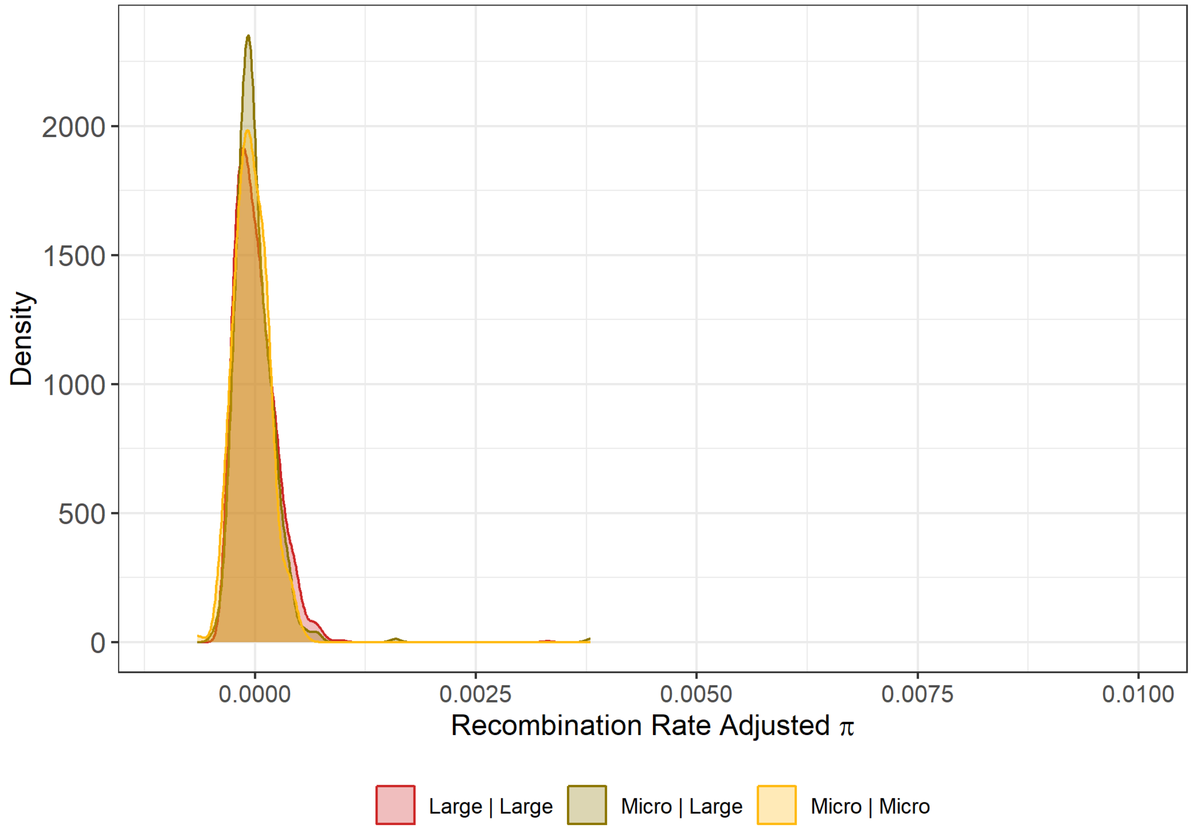


Supplementary Figure 14: Densities of residual nucleotide diversity, $\pi$, across 100KB windows by chromosomal history (Past | Current) in gyrfalcons after regressing on recombination rate.

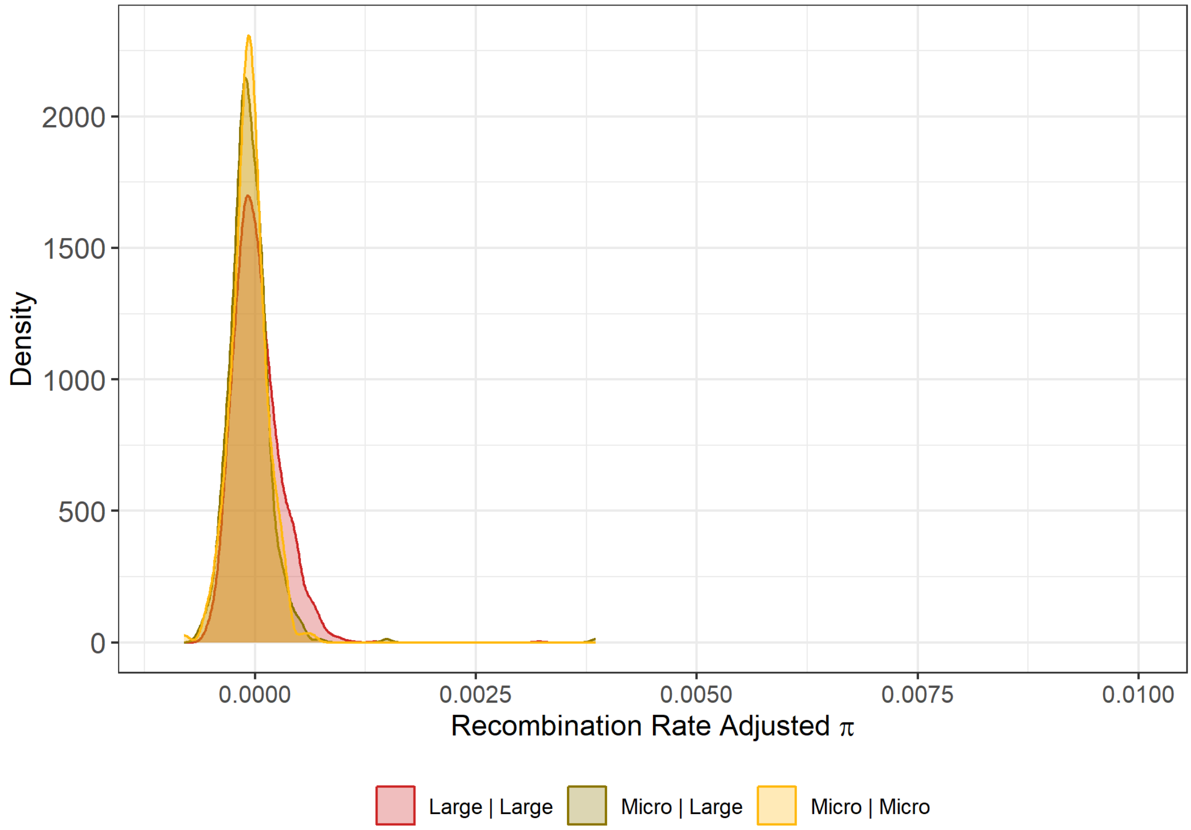


Supplementary Figure 15: Densities of residual nucleotide diversity, π, across 100KB windows by chromosomal history (Past | Current) in saker falcons after regressing on recombination rate.

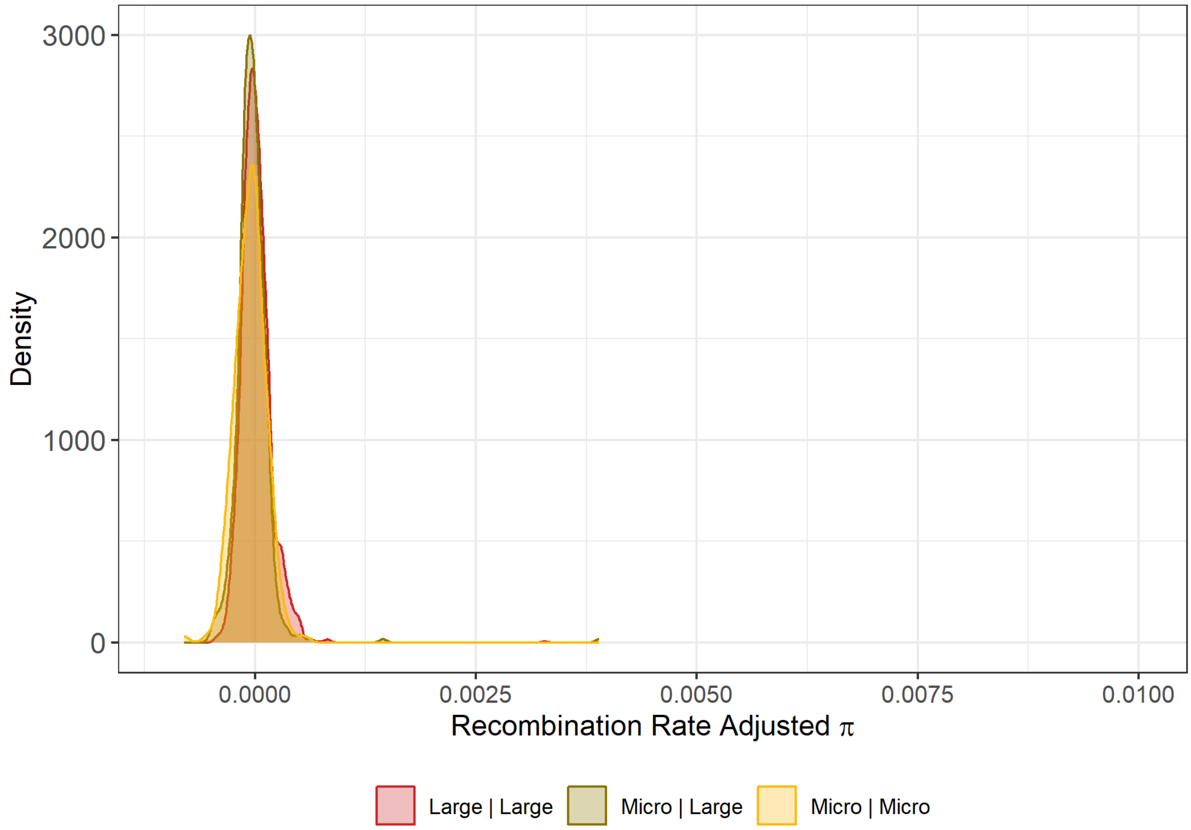


Supplementary Figure 16: Densities of residual nucleotide diversity, $\pi$, across 100KB windows by chromosomal history (Past | Current) in peregrine falcons after regressing on recombination rate.

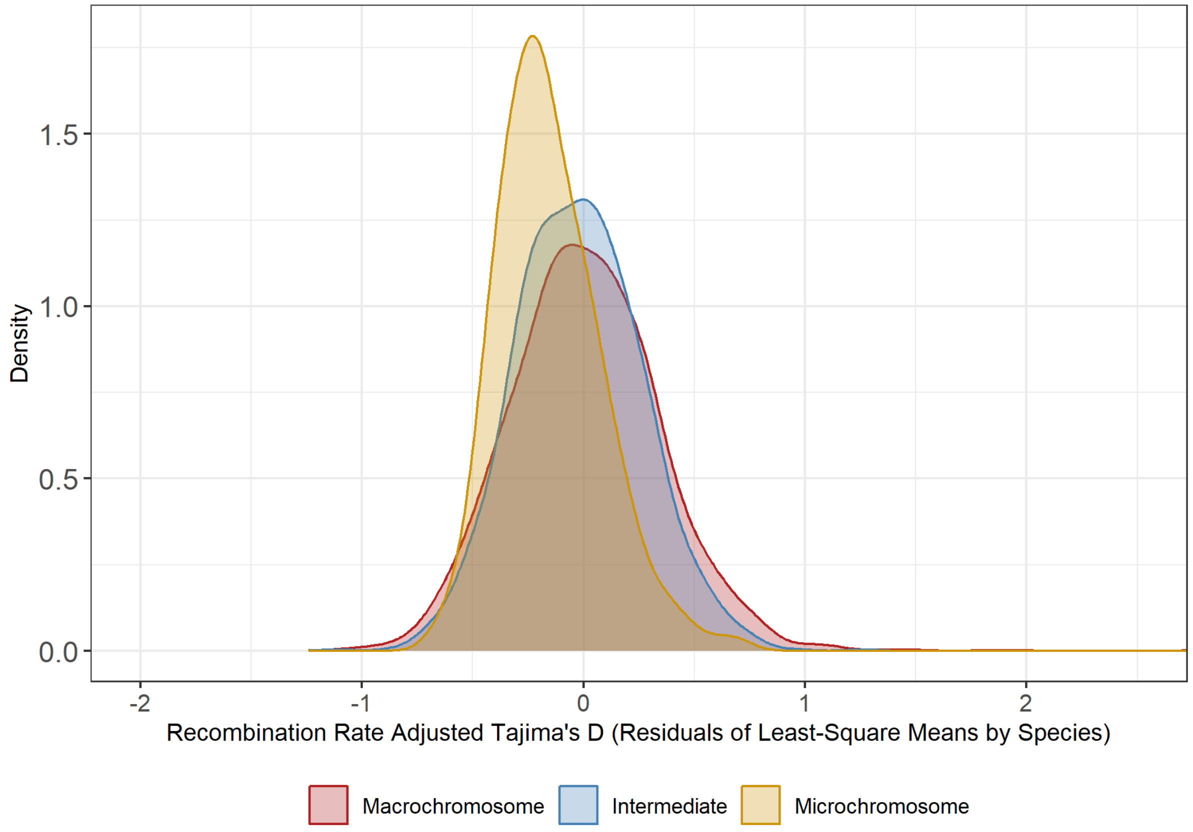


Supplementary Figure 17: Densities of residual Tajima's D across 100KB windows by current chromosome type after regressing first on species (least square means) then on recombination rate.

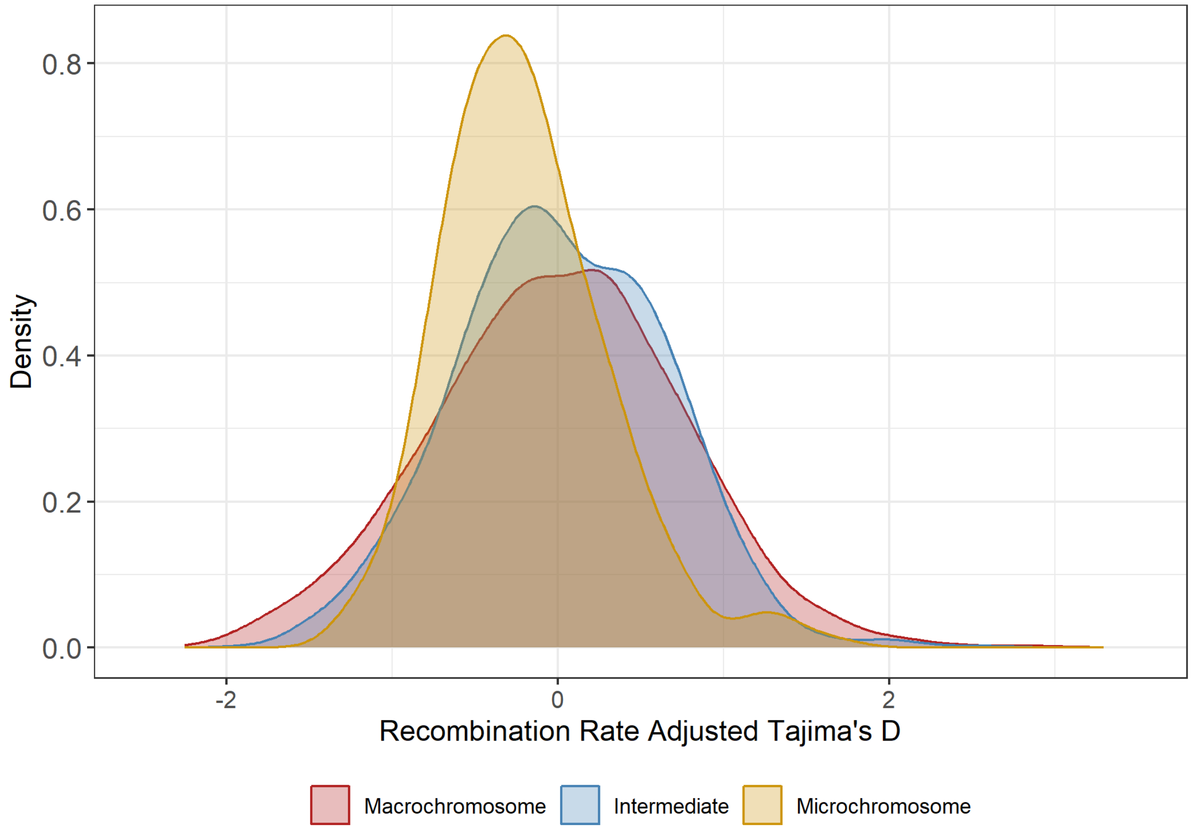


Supplementary Figure 18: Densities of Tajima's D across 100KB windows by current chromosome type in gyrfalcons after regressing on recombination rate.

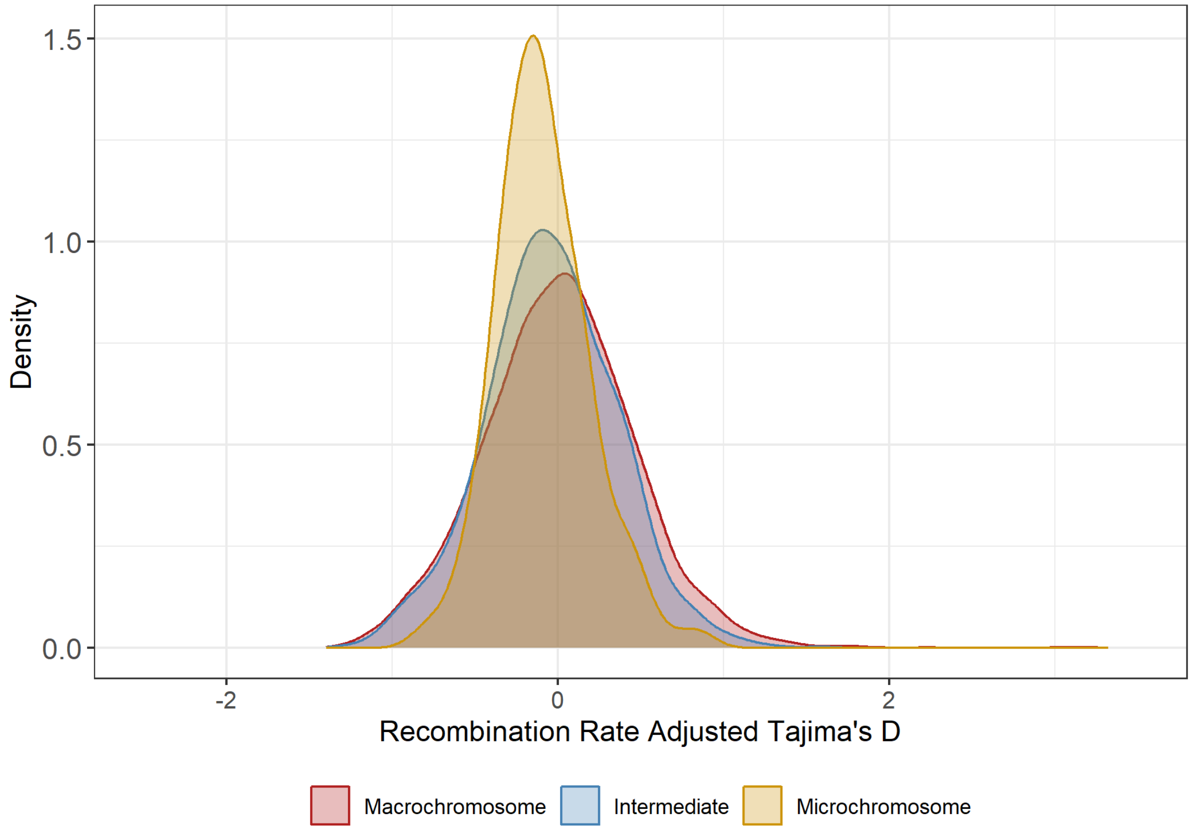


Supplementary Figure 19: Densities of Tajima's D across 100KB windows by current chromosome type in saker falcons after regressing on recombination rate.

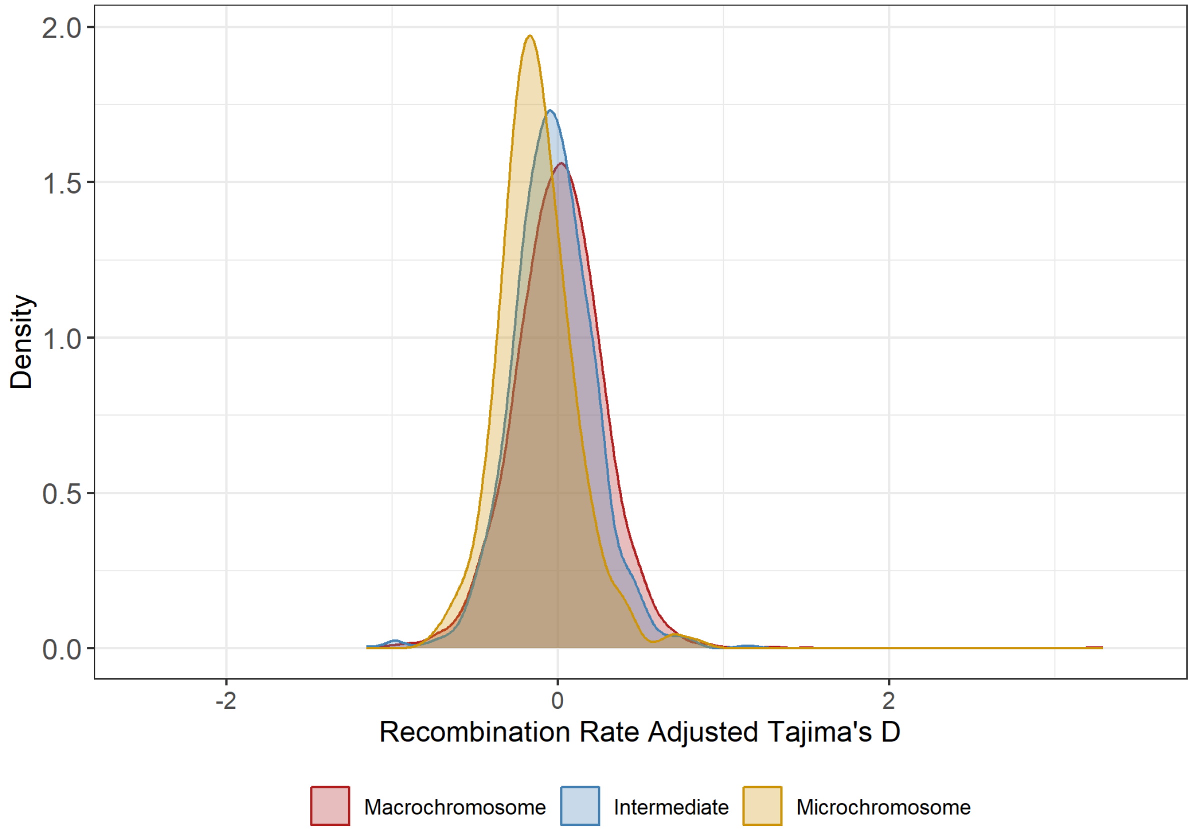


Supplementary Figure 20: Densities of Tajima's D across 100KB windows by current chromosome type in peregrine falcons after regressing on recombination rate.

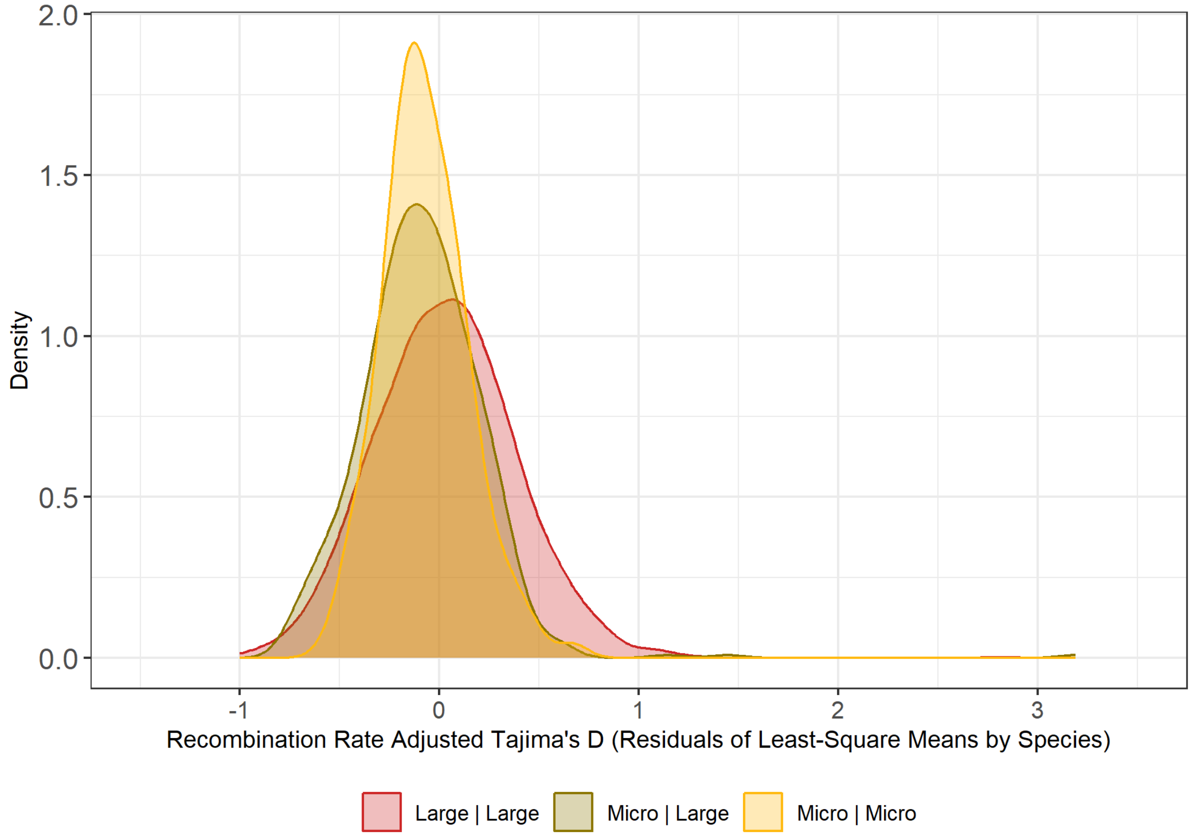


Supplementary Figure 21: Densities of residual Tajima's D across 100KB windows by chromosomal history (Past | Current) after regressing first on species (least square means) then on recombination rate.

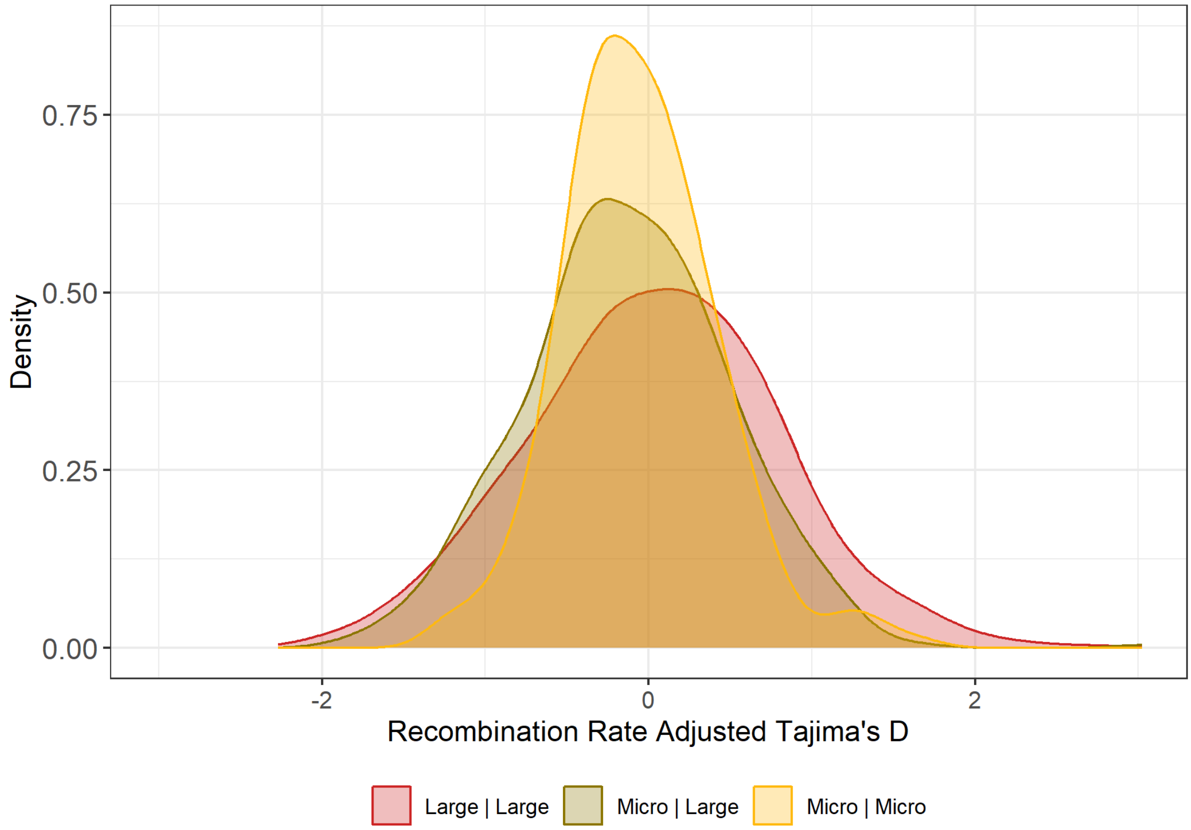


Supplementary Figure 22: Densities of residual Tajima's D across 100KB windows by chromosomal history (Past | Current) in gyrfalcons after regressing on recombination rate.

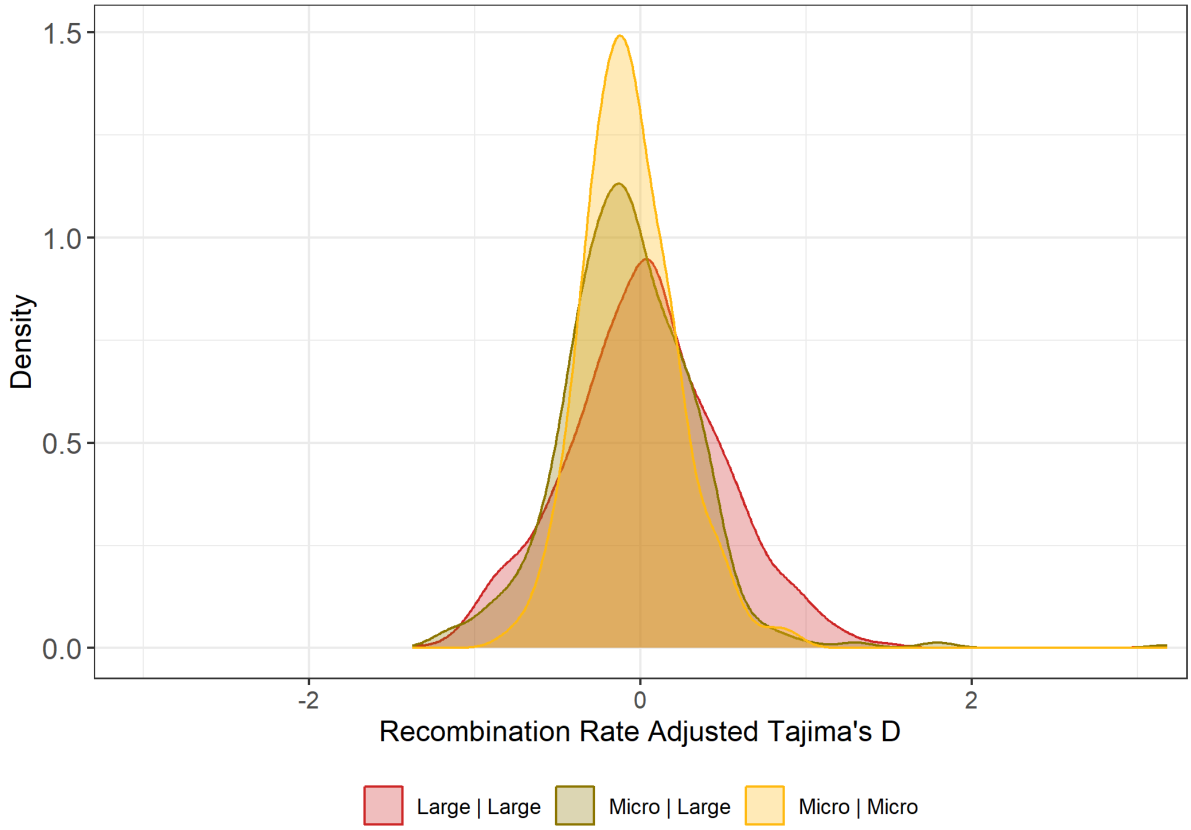


Supplementary Figure 23: Densities of residual Tajima’s D across 100KB windows by chromosomal history (Past | Current) in saker falcons after regressing on recombination rate.

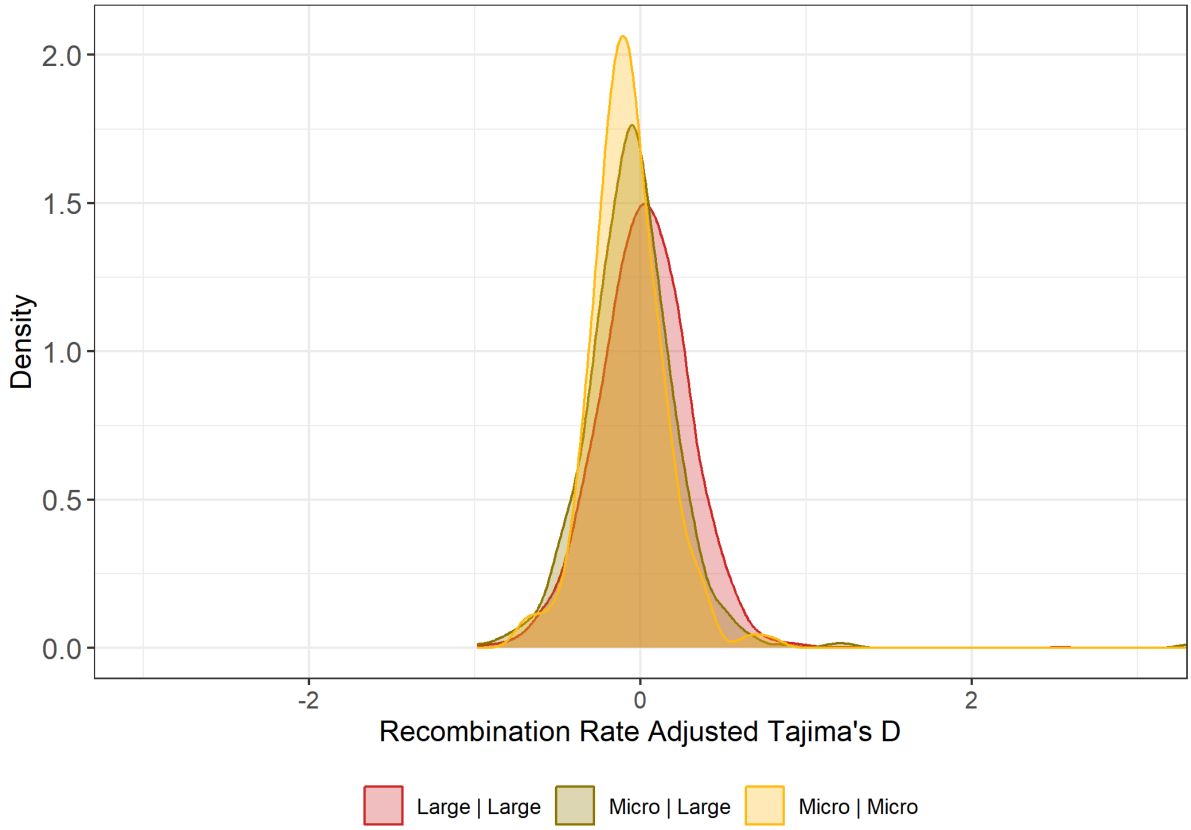


Supplementary Figure 24: Densities of residual Tajima’s D across 100KB windows by chromosomal history (Past | Current) in peregrine falcons after regressing on recombination rate.
